# CryoWriter: A Robotic Solution for Improved Cryo-EM Grid Preparation

**DOI:** 10.1101/2025.11.01.686008

**Authors:** KV Chinmaya, Babatunde Ekundayo, Marta Di Fabrizio, Inayathulla Mohammed, Julika Radecke, Henning Stahlberg, Massimo Kube

**Author notes:** UK Dementia Research Institute, Unit 3A, 338 Euston Road, London NW1 3BT, UK.

## Abstract

Cryo-electron microscopy (cryo-EM) structure determination relies on preparing thin, vitreous films of sample solution on EM grids. Cryo-EM is a mature technology, but preparing the grids remains a major bottleneck. Here, we evaluate the cryoWriter, a blotting-free, microfluidic grid-preparation robot that writes nanoliter volumes onto EM grids in a controlled environment. Using capillary-writing in spiral or line patterns, we prepared high-quality grids from minimal sample volumes and obtained near-atomic reconstructions for test specimens, including TMV, apoferritin, and the membrane protein TRPM4. We further demonstrate programmable deposition modes, such as writing the sample twice to boost particle density, or two-line writing for on-grid mixing to visualize time-resolved protein–ligand binding. In a challenging case (NrS-1 DNA polymerase), the cryoWriter grids exhibited reduced orientation bias relative to conventional blotting, enabling a more isotropic reconstruction. These results show that the cryoWriter provides a versatile platform for reproducible low volume cryo-EM grid preparation and for on-grid biochemical workflows.

## Introduction

Cryo-electron microscopy (cryo-EM) has evolved significantly since its introduction in the 1980s. Jacques Dubochet and colleagues had discovered that water could be quick-frozen by plunging into liquid ethane that was kept at its freezing temperature by liquid nitrogen. Quick-frozen water presented itself as an amorphous, highly viscous state of ice. This vitrified water maintains the native state of proteins and biological tissue that could then be imaged by cryo-EM ^1,2^. This sample preparation method has revolutionized structural biology by enabling the visualization of biomolecules at near-atomic resolution without the need for crystallisation ^3,4^

In its early days, vitrification for cryo-EM studies was performed using manual plungers. Typically 3 microliters of the sample solution are applied onto a grid, and 99.99% of that sample are blotted into a filter paper by hand, with only 0.01% of the remaining sample being quick-frozen by dropping the grid into liquid ethane using a simple, manually operated device. While this approach was groundbreaking, it was prone to inconsistencies in ice thickness, blotting time, and sample distribution, often resulting in suboptimal data ^5^. To address these challenges, automated systems began to emerge in the early 2000s. These included the Vitrobot (Thermo Fisher Scientific, TFS) and GP (Leica) grid plungers, which introduced precise environmental control, including temperature and humidity regulation, along with programmable blotting and plunging cycles, significantly improving reproducibility. Recent innovations have further automated the process, integrating advanced robotics for tasks such as grid handling, sample deposition, and vitrification, so that grids can be prepared with minimal human intervention. Notable examples are the Chameleon (SPT Labtech) ^6^, the VitroJet (CryoSol) ^7^, or the EasyGrid systems ^8^. Nevertheless, most approaches still require microliters of sample volume, sometimes at rather high protein concentration, of which only a minuscule fraction of a few picoliters ends up on the prepared cryo-EM grid.

The cryoWrite AG (Basel, Switzerland), has developed a robotic grid preparation device, called cryoWriter, that is based on a microfluidic approach to grid preparation. The cryoWriter almost fully automates grid handling, allowing to prepare cryo-EM grids from as little as 5 to 10 nanoliters of sample volume. The cryoWriter eliminates grid blotting by employing a micro-capillary system to precisely dispense nanoliter volumes of protein samples onto a 3 mm diameter cryo-EM grid that has been rendered hydrophilic by glow discharge in air. The concept of the cryoWriter is based on microfluidic strategies that were developed by the team of Thomas Braun at the University of Basel ^9–12^. The implementation in the cryoWriter combines the microfluidic handling of the sample with automated grid storage, grid glow discharging, sample “writing” onto the grid, grid plunging and storing under liquid nitrogen and re-warming of the plunging tweezer, so that several grids can be prepared and frozen in a fully automated series. Based on the approaches by the Braun team ^13^, the cryoWriter dispenses sample through a process of capillary writing onto the grid, but without re-uptake of sample as previously done by the Braun team.

The sample is dispensed from the capillary using a spiral or line writing, with nanoliters per second being dispensed at high precision, By optimizing parameters such as the speed of the capillary movement, the distance between the capillary and the grid, and the speed of the sample dispensing, a precise control over the sample layer thickness can be achieved. We found this approach to enable the reproducible production of high-quality grids.

We here provide a systematic analysis of the cryoWriter, its advantages and difficulties, and present novel application methods that allow sample mixing on the grid and give access to time-resolved cryo-EM investigations.

## Results

### The cryoWriter as robotic sample handling and grid vitrification station

The cryoWriter is a fully automated cryo-EM sample preparation robot housed in a 150 × 200 × 140 cm³ enclosure (**Fig. 1a**). Its central element is a microfluidic glass capillary that aspirates nanoliter volumes of sample and deposits them in precise patterns onto cryo-EM grids. Grid handling is performed by a robotic tweezer (“gripper”), which transfers grids between the storage box, the integrated glow discharge unit, the writing platform (“launchpad”), the plunge freezer, and liquid-nitrogen storage. An integrated light microscope allows observation of samples, including cells and tissues, directly within the cryoWriter.

**Fig. 1.**
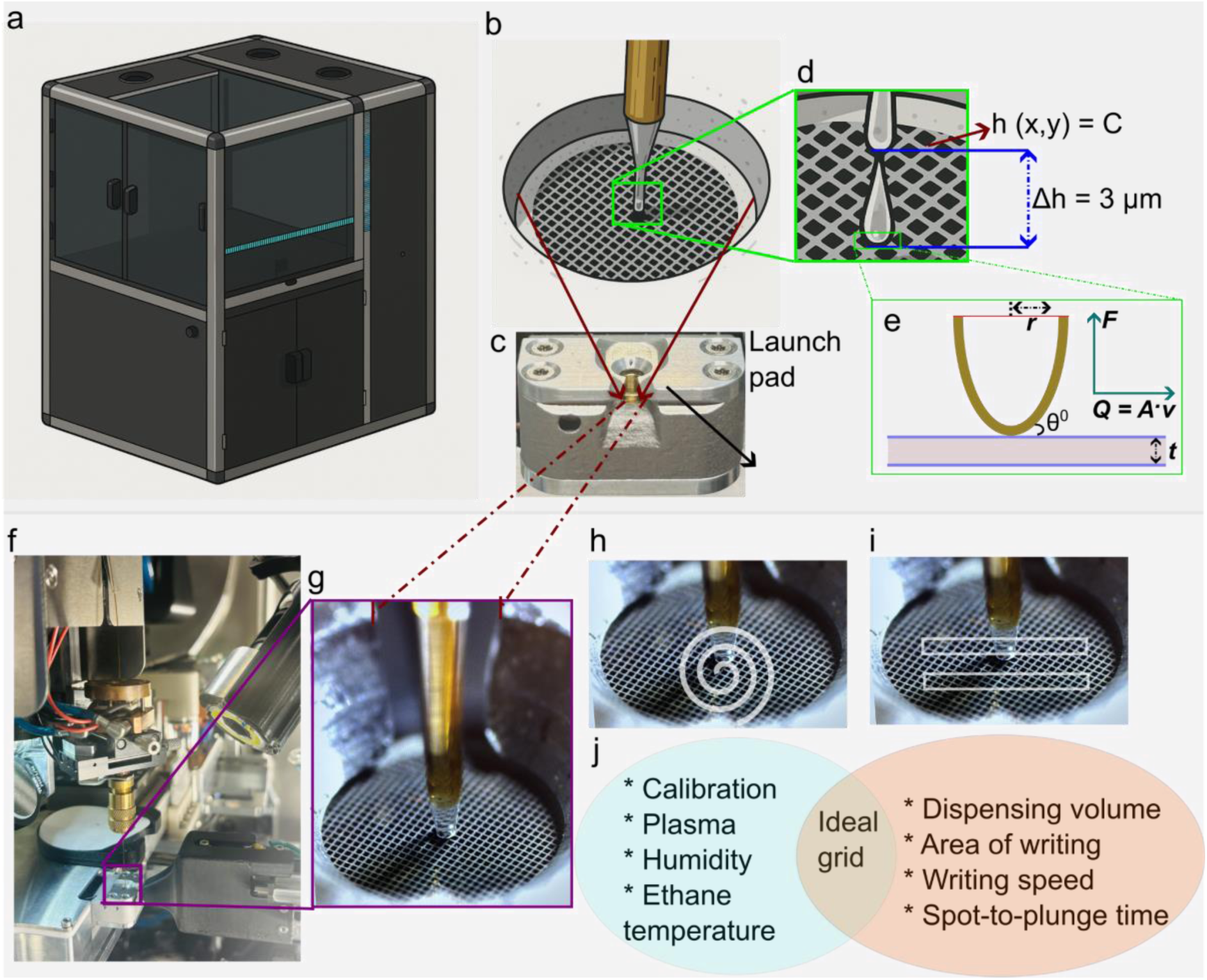
Internal and external architecture of the cryoWriter system and sample preparation workflow. **a** Schematic representation of the external architecture of the cryoWriter instrument. **b** Diagram illustrating the cryoWriter sample writing mechanism. **c** The launchpad used for grid clamping and cooling to dewpoint temperature. **d** Schematic showing the optimal distance between the capillary tip and the cryo-EM grid during sample deposition. **e** Diagram illustrating capillary-based sample writing and the effects of liquid flow dynamics during deposition. **f** The freezing setup showing both the capillary robot and the tweezer robot in operation. **g** Grid positioned on the launchpad with the capillary robot precisely aligned for sample writing. **h** Spiral writing pattern illustrating sample deposition on the grid surface. **i** Illustration of single-line and double-line writing patterns used for sample deposition. **j** Schematic representation of optimal grid freezing parameters in the cryoWriter workflow.

The workflow proceeds as follows: the gripper retrieves a grid from a box and places it on the glow discharge platform. A 3-cm-diameter bell cover seals the platform, the chamber is evacuated, and the grid is glow discharged. After venting, the gripper transfers the activated grid to the launchpad. A microfluidic pipette then aspirates a few nanoliters of sample and deposits them onto the grid in programmable patterns. Within 200 ms, the gripper plunges the prepared grid into liquid ethane at −183 °C to vitrify the sample. The vitrified grid is finally transferred into a storage box under liquid nitrogen. The full cycle from retrieval to storage takes three to four minutes and runs fully automatically (**Suppl. Video 1)**.

The key method for sample application is capillary writing (**Fig. 1b**). Writing parameters such as the diameter of the employed capillary, the capillary writing speed, writing area, and dispense rate, can be adjusted to control sample deposition and ice thickness. A precise distance between the capillary and the grid, typically of three micrometers, together with precise robotic movement of the capillary, ensures uniform and controlled deposition of the sample (**Fig. 1c**). The capillary is driven by a motorized pipette that can reproducibly aspirate and dispense nL volumes. This allows the capillary to dispense minimal sample volumes (3 to 6 nL) at continuous and homogeneous speeds over a few seconds, thus eliminating the need for blotting. The grid is held flat in the *launchpad*. The grid’s flatness is defined as *h*(*x,y*) (**Fig. 1d**), with *h* being the height at position (*x,y*), which should remain at a constant value *C* throughout the entire grid surface if the grid is flat and not damaged. The optimal target value of *C* depends on the grid type, the sample buffer, and the local temperature. Because thermal expansion can cause the grid, launchpad, and capillary to expand or contract, the system is calibrated at a specific temperature to maintain a consistent height C during sample preparation. The ideal grid position for writing is shown in **Fig. 1f** and **g**.

The cryoWriter is controlled by the cryoWrite OS operating system. The software supports two default writing modes, a *spiral* and a *line* mode, as shown in schematic diagrams in **Fig. 1h** and **Fig. 1i**. For the demonstration, the spiral deposition of the protein sample on the Ultrafoil grid is shown in **Suppl. Video 2**. The cryoWriter maintains an elevated relative humidity in the enclosure at a user-defined value, typically 70%, and the cryoWrite OS calculates the corresponding dewpoint temperature, with the option for the user to define a small temperature offset if needed. Typically the grid is kept slightly (1-2°C) above the dew point temperature to prevent water condensation or allow for controlled evaporation. Additionally, the cryoWriter features a temperature-controlled sample storage block (nano-incubator), whose temperature can be adjusted between -2 °C and +60 °C to meet the requirements of the sample. This allows temporary storage of biological samples, e.g., provided in small Eppendorf cups. The grid is eventually plunged into a metal cup filled with liquid ethane, which, prior to grid preparation, has to be prepared by the operator by cooling its surroundings with liquid nitrogen and blowing ethane gas into the cooled cup for liquefaction of the ethane. For grid plunging, the temperature of the ethane cup can then be adjusted precisely with the help of a built-in thermostat. This allows keeping the ethane liquid for prolonged times at a desired temperature, typically between -180 to -183 °C, so that rapid vitrification of the sample can be achieved.

### High-resolution cryo-EM data from cryoWriter grids

Initial optimization efforts for sample preparation revealed issues such as excessive evaporation, dryness, crystalline ice formation, and overly thick ice conditions unsuitable for data collection, as shown in the **Suppl. Fig. S1**. In order to obtain highly reproducible writing of homogenous and thin vitreous ice, several parameters, such as the writing speed, capillary distance, and writing volume, were optimized, which enabled spiral and line writing (**Fig. 2** and 3, **top panels**). During the initial optimization, constant writing start and end speeds (2 mm/s and 8 mm/s, respectively) were used, which resulted in a mixture of thick and thin ice. Modulating the writing speed enabled the generation of a controlled gradient of amorphous ice. The capillary–grid distance was maintained at a constant value of 3 µm throughout the writing process to ensure uniform sample distribution over the defined surface area of the grid. The chamber humidity was maintained at 60–70% to prevent sample drying. When the humidity dropped below 60%, rapid evaporation of the thin liquid film was observed, leading to dehydration and formation of crystalline ice (**Suppl. Fig. S1**). At slower writing speeds (2 mm/s), dispensing larger volumes of sample (5–8 nL) frequently resulted in thicker ice, particularly during line writing.

**Fig. 2.**
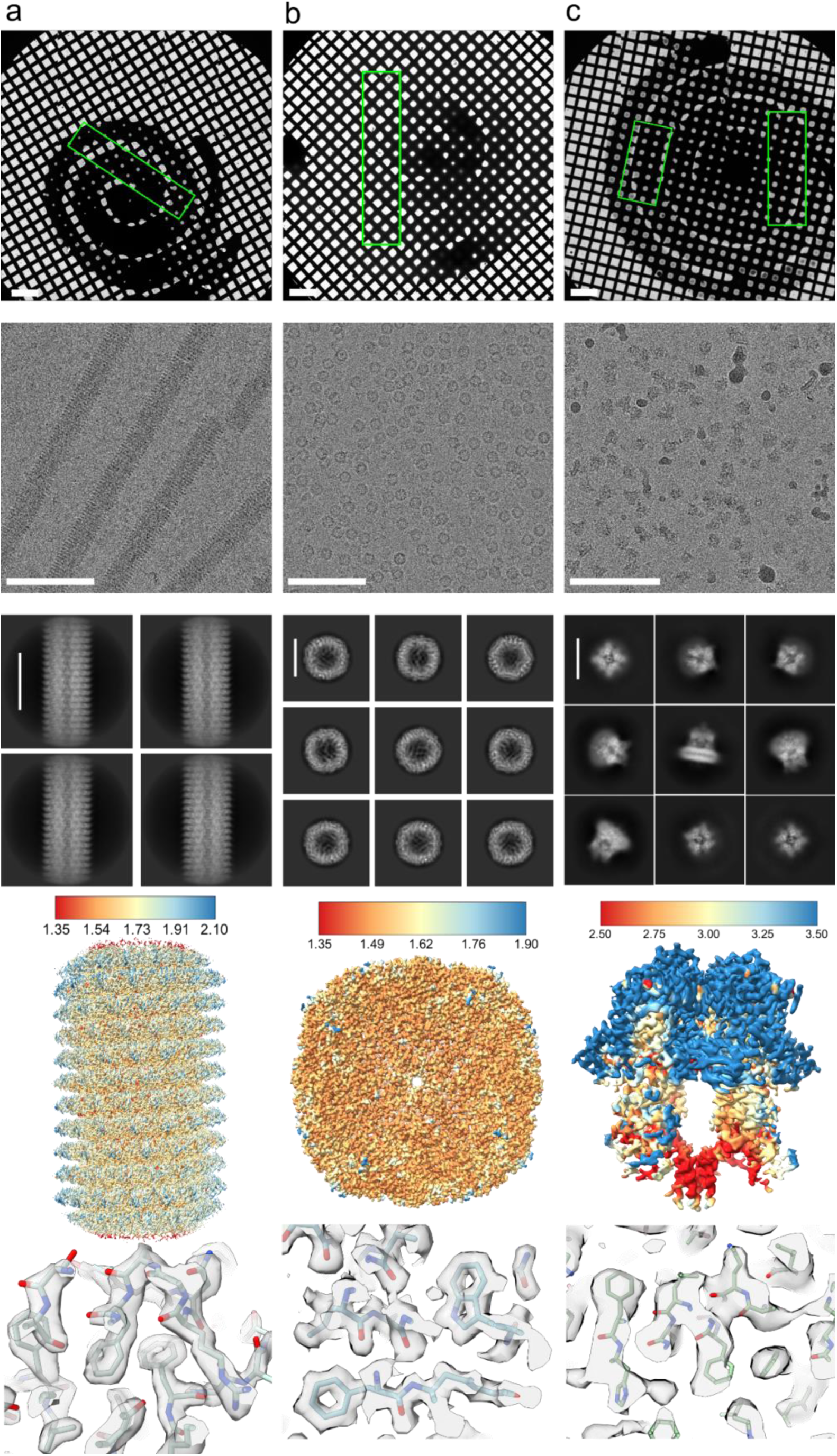
High-resolution single particle reconstruction of various types of protein samples prepared using the cryoWriter. **a** Tobacco mosaic virus (TMV), **b** Horse spleen apoferritin (apoF), **c** Transient receptor potential melastatin 4 (TRPM4) channel. For each sample from top to bottom: an atlas image, indicating the areas where data were collected by green boxes (scale bar = 200 μm); Representative cryo-EM micrograph (scale bars from left to right = 60 nm, 80 nm, 100 nm); 2D class averages of picked particles (scale bars from left to right: 19 nm, 10 nm 16 nm); 3D reconstructions with local resolution shown by the color scales (in Å); Representative regions in the 3D reconstructions, with fitted atomic protein models.

**Fig. 3.**
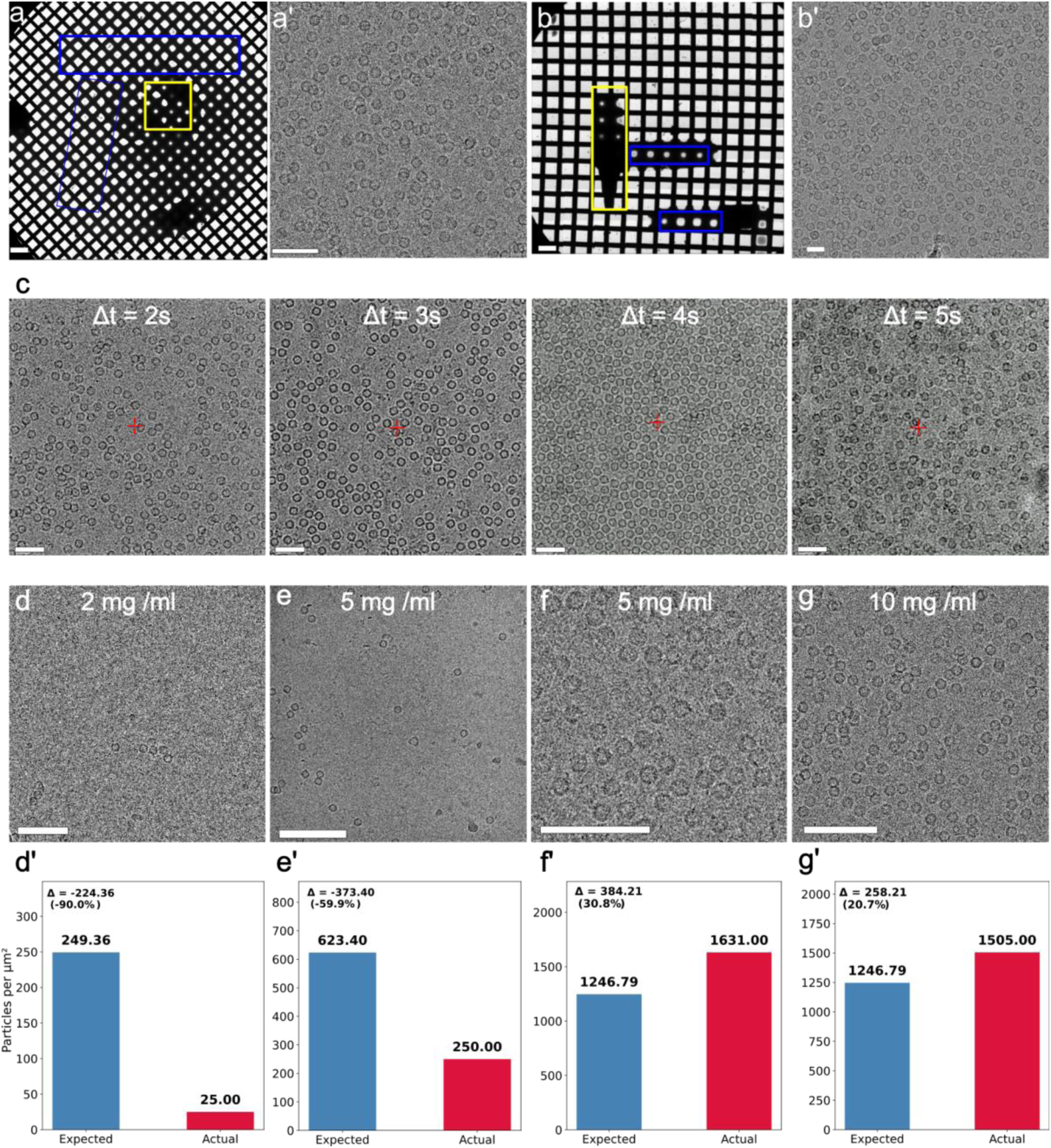
Capillary writing of the apoferritin sample and its distribution in an electron micrograph. **a** Low-magnification image (atlas) of the apoferritin sample demonstrating uniform, optimally thin ice, suitable for high-resolution structural analysis (scale bar = 200 μm), followed by an electron micrograph showing uniform ice and particle distribution at a concentration of 10 mg/mL in **a’** (scale bar = 50 nm). Yellow boxes: indicate the thick ice region. Blue boxes: indicate the thinner ice region suitable for data collection. **b** Low-magnification image (atlas) of line writing of the apoferritin sample, demonstrating uniform, optimally thin ice (scale bar = 200 μm), followed by an electron micrograph showing particle distribution at a concentration of 10 mg/mL **b’** (scale bar = 20 nm). **c** Capillary writing of the apoferritin sample and corresponding electron micrograph showing particle distribution, performed with two times writing at a concentration of 5 mg/mL and varying waiting times (2 to 5 seconds) before plunge freezing (scale bar = 40 nm). **d** Theoretical and experimental apoferritin particle distributions at different concentrations, assuming an ice thickness of 100 nm. **d–g** Electron micrographs of apoferritin particle distribution at varying concentrations (scale bar from left to right = 50, 20, 100, and 80 nm). **d’–g’** Normalized graphs of apoferritin distribution per μm² in the corresponding electron micrographs, comparing theoretical predictions with the experimental mean particle distribution.

To validate the cryoWriter sample preparation method, several different protein systems were used as test samples, including tobacco mosaic virus (TMV), which is widely used as benchmark for helical samples, horse spleen apoferritin (apoF), and transient receptor potential melastatin 4 (TRPM4), a calcium-activated monovalent cation channel membrane protein, associated with various genetic and cardiovascular disorders ^14,15^. Cryo-EM reconstructions of these samples prepared with the cryoWriter are shown in **Fig. 2**. We optimized the operation protocols, which now allow to routinely produce consistent, high-resolution yielding grids.

Grids from tobacco mosaic virus (TMV) were prepared with a spiral writing pattern at a concentration of 20 mg/ml, using a starting speed of 2 mm/s in the center of the spiral and increasing to 8 mm/s at the outer edge of the spiral, while dispensing sample at a constant dispensing speed of 5.5 nL in 3 seconds onto the cryo-EM grid. A gradient in ice thickness within the written spiral trace is visible, with thinner ice towards the outer edges of the spiral. A dataset of 5,824 movies was recorded from thinner ice locations, which revealed intact helical TMV particles, which allowed the determination of the TMV structure from 95,222 helical segments at a resolution of 1.83 Å (**Fig. 2a**, **Suppl. Fig. S2a**, and **Suppl. Table 4**).

Grids from horse spleen apoferritin (apoF) were equally prepared with a spiral writing pattern at a concentration of 10 mg/ml, dispensing 5.5 nl of sample within 2 seconds at a writing speed ranging from 2 mm/s to 8 mm/s. A final 3D reconstruction at 1.68 Å resolution was obtained from 469,137 particles (**Fig. 2b**, **Suppl. Fig. S2b**, and **Suppl. Table 4**).

For the transient receptor potential melastatin 4 (TRPM4) channel protein present in detergent (0.02% GDN), cryoWriter grids prepared with a spiral writing pattern at a concentration of 6 mg/ml, dispensing 4.15 nl of sample within 2 seconds at a writing speed ranging from 3 mm/s to 8 mm/s, allowed the reconstruction of the membrane protein structure in detergent at 3.03 Å resolution from 69,878 particles, refined in C4 symmetry (**Fig. 2c**, **Suppl. Fig. S2c**, and **Suppl. Table 4**).

ApoF was also prepared for cryo-EM using the cryoWriter with a line-writing approach, which resulted in a reduced usable grid area. A total of 4,923 micrographs were collected. Grids were prepared by two consecutive line-writing protocols at a speed of 8 mm/s, depositing approximately between 2–3 nL of sample per grid. This enabled reconstruction of the ApoF structure at 2.02 Å resolution from 230,450 particles refined in O symmetry **(Suppl. Fig. S3).**

The FSC resolution estimations for all samples (**Suppl. Fig. S2**) demonstrate that the cryoWriter can be used to prepare high-quality cryo-EM grids from a wide range of protein samples, including helical viruses, isolated single particles, and detergent-solubilized membrane protein particles.

### Sample deposition by capillary writing for single particle cryo-EM

The concentration of the sample needed for the production of optimal single-particle data acquisition grids was slightly higher than needed for some other semi-automated grid preparation techniques. The quantitative comparison of particle yield for final 3D reconstruction and concentration between cryoWriter and conventional plunge freezing is presented in **Suppl. Table 1**. To further investigate this, we explored capillary-writing deposition of the sample at varying waiting times and concentrations. Apoferritin (apoF) was used as a test sample, as it is a widely used standard sample in cryo-EM and allows reaching 1.09 Å resolution ^16^. We prepared grids with different approaches, including spiral writing (**Fig. 3a**) and line writing (**Fig. 3b**) at high concentrations (10 mg/ml), using different spot-to-plunge times (**Fig. 3c**), and also employing a “multiple writing” mode (**Fig. 3d**). The particle density of the sample on the grids can be increased by allowing for different absorption and spot-to-plunge times prior to freezing (**Fig. 3c**). In contrast to the standard protocol available in the software, where the grid is plunged almost instantaneously after writing, the grid with the applied sample can be kept on the *launchpad* for a slightly longer duration, allowing sample equilibration and controlled sample evaporation. When slightly more sample volume was applied to the grid and the grid was maintained in the *launchpad* at a temperature slightly above the sample’s dew point, the water in the sample slowly evaporated in a controlled manner, causing an up-concentration of the sample on the grid. Using this approach, we created grids with increased particle densities (**Fig. 3c** and **Suppl. Table 2**). However, after longer drain times, the sample showed signs of deterioration (**Fig. 3c, last image**).

We also performed experiments in which we applied the sample twice onto the same grid, with the second writing process starting immediately after the first. The subsequent time interval between the last sample application and grid vitrification (spot-to-plunge delay) was also systematically varied between 1 and 5 seconds to allow for different degrees of concentration. Representative micrographs of apoferritin at delay times between 2 and 5 seconds are shown in **Fig. 3c**. For this sample, we found that a suitable spot-to-plunge time delay window was 2 to 4 seconds, within which particle integrity was preserved. Delays exceeding 4 seconds resulted in signs of particle denaturation.The deterioration observed in both cases is likely due to increased evaporation with a corresponding increase in salt concentration due to the longer time before plunging, combined with a prolonged exposure of the protein samples to the air–water interface.

To further characterize the effect of initial sample concentration on particle density across the grid, we prepared grids from samples at varying protein concentrations. For single-spiral writing with 2 mg/mL, 5 mg/mL, and 10 mg/mL samples, the mean observed number of particles per µm^2^ was 26, 250, and 1505, respectively. Experimental details are provided in **Suppl. Table 3**. The recorded cryo-EM images of these grids were analyzed for observable particle density (**Fig. 3d–g**) and compared with the theoretically expected particle density (**Suppl. Text 2**), assuming a constant ice thickness of 100 nm (**Fig. 3d’–g’**). At an apoF concentration of 5 mg/mL, the observed particle density increased from 250 particles/μm² under single writing conditions to 1631 particles/μm² with double writing. Notably, at a lower concentration of 2 mg/mL, the experimentally observed particle density was more than tenfold lower than the theoretical estimation. This non-linear relation between observed and applied particles is in agreement with the hypothesis that particles first have to cover the grid and carbon film surfaces, after which remaining additional particles fill the ice holes. These results indicate that applying double writing at lower protein concentrations can substantially increase particle density in cryo-EM micrographs, thereby facilitating faster data acquisition and reducing the total amount of data required.

Given the dependence of the particle density and waiting time, we investigated the effect of the air-water interface on particle distribution across the gradient of ice thickness. Previous studies ^17,18^ have shown that protein molecules could adhere to the air-water interface and also be denatured at this interface, influencing the particle distribution across the ice. To characterize the ice gradient and examine the air-water interface across the grid, we used apoF samples at 5 mg/mL and performed one or two successive spiral writing applications onto the grid. The capillary writing was done with a starting speed of 2 mm/s, gradually increasing to an end speed of 8 mm/s, after which the ice thickness and particle distribution could be analyzed using cryo-electron tomography (cryo-ET) as shown in **Suppl. Text 2 & 3** and **Fig. 4**.

**Fig. 4.**
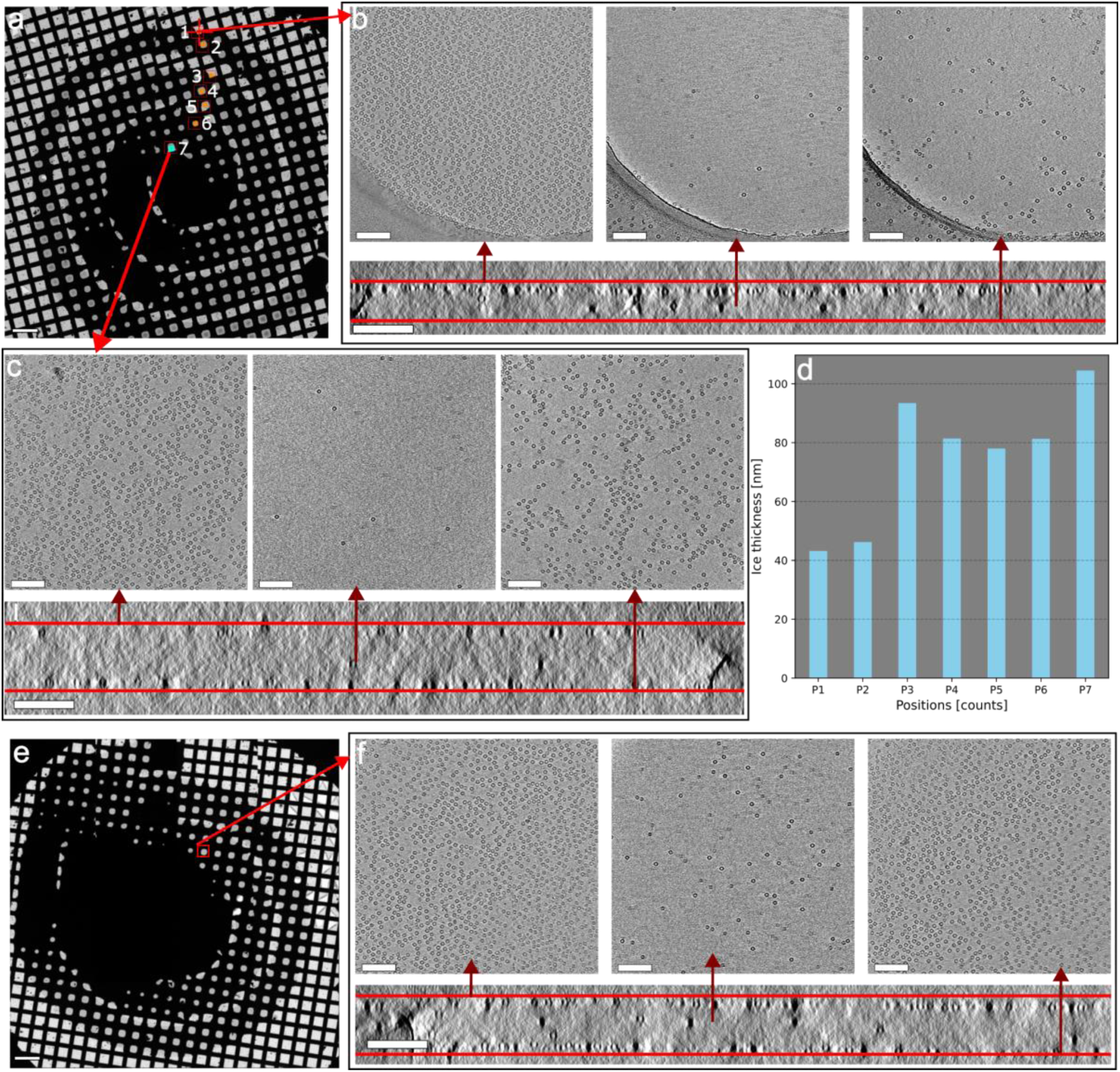
Ice thickness measurement on a cryoWriter grid using cryo-ET. **a** Low-magnification image of apoF frozen at 5 mg/mL. Position numbers show the regions from the end of writing to the start of writing. **b** Tomographic reconstruction of the sample in (a), shown at three different height levels (top, center, bottom) across the 3D reconstruction. The top layer was also on the top when writing the sample onto the grid with the cryoWriter, which applied the sample with the capillary from the top to the surface. ApoF particles can be seen mostly at the upper air-water interface (AWI), shown in the first of the three panels. **c** Tomographic reconstruction and apoF distribution across the depth of the ice layer of position 7. **d** Measured ice thickness for all positions shown in (a). **e** Low magnification image of apoF at 5 mg/mL and written twice. **f** Tomographic reconstructions and apoF distributions across the depth of a vitreous sample showing the AWI and apoF at the ice layer.

To better characterize the differences in ice thickness and particle distribution, we collected cryo-ET tilt series of an apoF grid (prepared with 5mg/ml concentration) at various locations (indicated by numbers in **Fig. 4a**) and reconstructed tomograms of these (**Fig. 4b,c,f**) to measure ice thickness. In all regions, we observed most particles at the air-water interfaces. In some grid regions, particles were found to be mostly adhering only to the one air-water interface on the side, from which the sample was written onto the grid. In these areas, only few particles were observed in the central region of the ice layer, and the opposing air-water interface had a significantly lower number of particles than the upper interface (**Fig. 4b**). The particle distribution within the volume is apparent in the cross-sectional view of the reconstructed tomogram volume (**Fig. 4b**). We collected five tomograms from different holes at position P1 in Fig. 4a, and the average ice thickness was determined to be 43.2 nm (**Fig. 4j**). We reconstructed a tomogram at position P7 (**Fig. 4c**), corresponding to the start of the writing path. At this position, we encountered a thicker ice layer of 105 nm (**Fig. 4c**). We quantitatively measured mean ice thickness at all positions from P1 to P7, as presented in (**Fig. 4d**). Here, high particle concentrations were found on both surfaces of the ice layer. Tomograms at intermediate positions P2 to P6 along the spiral writing path are discussed in **Suppl. Fig. S4**, each writing showing that particles were mostly located on the air-water interface and only a small fraction in the centre of the ice layer, as also observed before by Noble et al. (2018) ^19^. Although increasing the writing speed compared to semi-automated approaches may enable the formation of a more favorable ice gradient, minimizing particle adhesion to the air-water interface, which can lead to sample denaturation and preferred orientation, remains a significant challenge, even when using the cryoWriter.

### Less preferred particle orientations in grids prepared with the cryoWriter

Orientation bias resulting from sample adhesion to the air-water interface is a major limitation to obtaining high-resolution 3D reconstructions of many macromolecular systems analyzed by single particle cryo-EM ^20–23^. The use of the cryoWriter could be a powerful alternative to mitigate the orientation bias of some samples in cryo-EM experiments since we are able to obtain a controlled gradient distribution in ice, using the cryoWriter, compared to semiautomated approaches.

The NrS-1 DNA polymerase, a DNA polymerase identified in the deep-sea vent phage NrS-1 ^24^, is a hexameric DNA-binding protein. Other similar molecules have been widely studied by cryo-EM. However, in our laboratory we found NrS-1 to be an especially challenging protein for cryo-EM grid preparation. When preparing grids with a TFS Vitrobot Mark IV, this protein showed strong preferential orientation, hindering high-resolution structure determination (**Fig. 5a**) and only resulting in a reconstruction with anisotropic resolution of 3.8 Å. The sampling compensation factor (SCF), a measure of the degree of preferred orientation, was 0.448. Instead, grids prepared with the cryoWriter showed these particles in more random orientations (**Fig. 5b**), which allowed the determination of the 3D structure at 3.2 Å resolution. Here, the more broadly distributed range of orientations allowed the determination of the structure at a more isotropic resolution. Here, SCF improved to 0.649. While with this specific protein we observed a strong advantage with the cryoWriter, other proteins may behave differently. The different behaviors of the particles in cryo-EM grids prepared with one or the other method are not well understood. However, we note that the observed particle orientation bias in grids prepared with the capillary writing mechanism in the cryoWriter was less pronounced than when preparing grids with a filter-paper blotting method. The cryoWriter is therefore an interesting alternative tool to reduce orientation bias for challenging samples.

**Fig. 5.**
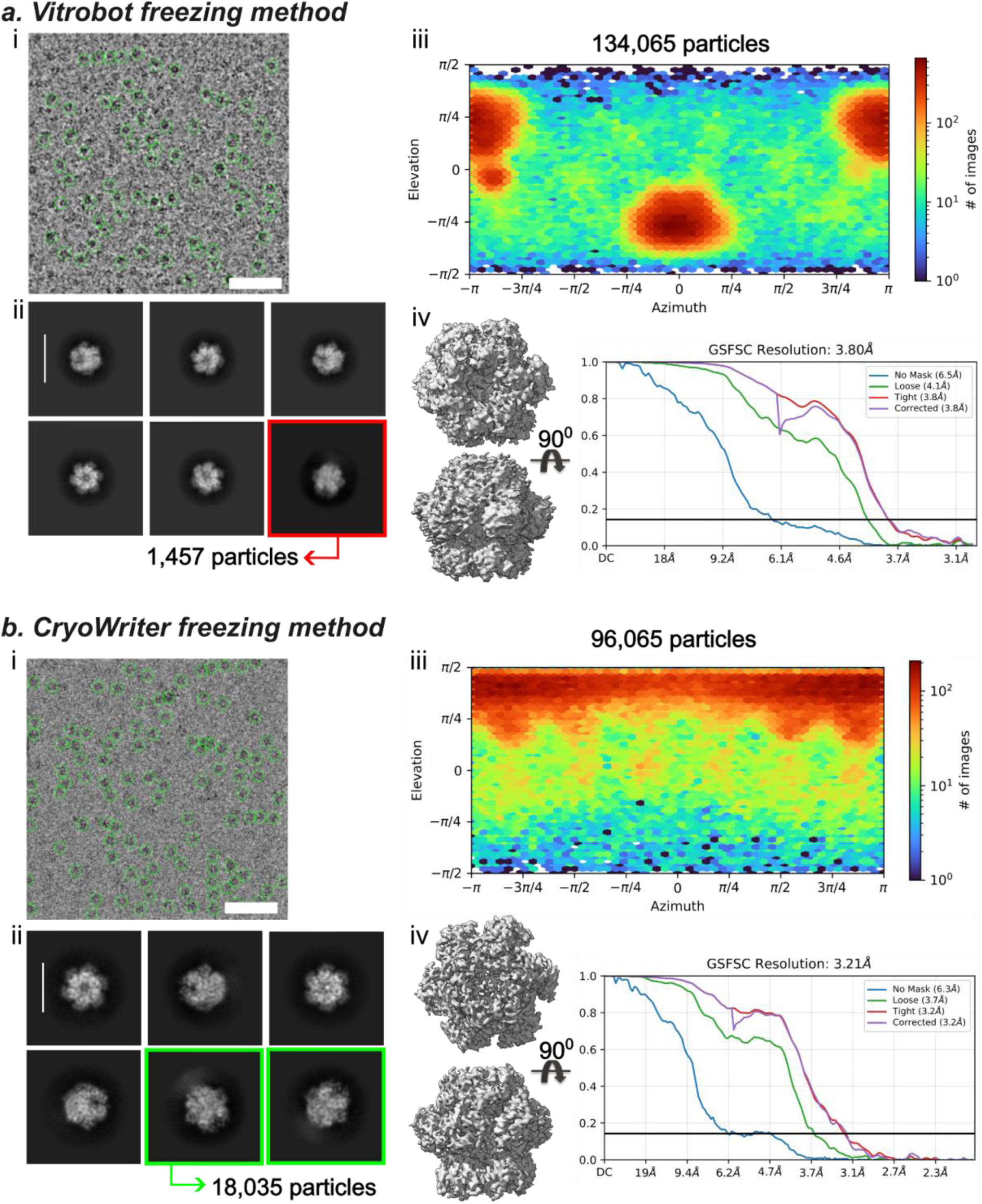
Comparison of the particle orientation in grids prepared with the TFS Vitrobot and cryoWriter, using Nrs-1 protein particles. **a** Vitrobot freezing method. i, Micrograph and particle distribution. ii 2D class averages (scale bar = 15 nm), with 1,457 side-view particles out of 134,065 total particles. iii Viewing direction distribution of particles from all angles with C1 symmetry applied. Side-views are mostly absent for this difficult protein. iv 3D reconstruction and FSC curve of the Nrs-1 frozen from the Vitrobot. **b** cryoWriter freezing method. i Micrograph and particle distribution. ii 2D class averages (scale bar = 11 nm) processed from the data set (scale bar = 15 nm), where 18,035 side view particles were found from 96,065 total particles used in 2D class averages. iii Viewing direction distribution of particles from all angles with C1 symmetry applied. iv 3D reconstruction and FSC curve of the Nrs-1 frozen with the cryoWriter.

### Protein interaction studies through simultaneous deposition of two different samples onto the same grid

Structure-based drug discovery (SBDD) is a critical strategy for the rational design and optimization of novel therapeutic agents ^25,26^. In cryo-EM–based drug screening and discovery, the ability to rapidly and reliably compare different sample states is essential for identifying ligand-induced conformational changes. Co-depositing two distinct samples onto the same grid, such as apo and ligand-bound forms of the same protein, allows for direct comparison under identical vitrification and imaging conditions, thereby improving screening efficiency, conserving sample material, and reducing dataset variability.

Deposition of multiple samples in a single writing step enables several additional applications for the cryoWriter. As a first approach, we wrote two different samples onto the same grid and studied the subsequent mixing of the two samples. This was done by sequentially aspirating two different samples into the same capillary, separated by a small air bubble, and writing both samples subsequently onto the same grid as two closely adjacent but not overlapping lines (**Suppl. Fig. 6**).

As a test sample, we deposited NrS-1 and apoF onto the same grid using the cryoWriter. NrS-1 protein was deposited at a concentration of 6 mg/mL as the first line, followed by apoF protein at 10 mg/mL concentration as a parallel line with a 300 μm gap separating them (**Fig. 6**). The writing speeds for both NrS-1 and apoF were maintained at 8 mm/s. A total of 4 nL of NrS-1 and 6 nL of apoF were aspirated, separated by a 2 nL air bubble. These two samples were then written in one U-shaped trajectory onto the same grid. This allowed recording images of NrS-1 alone in one zone, apoF in another zone, and a mix of both samples was found in the intermediate zone between the two written lines (**Fig. 6a-f**). In micrographs recorded at the center between the two writing lines, the particle ratio between NrS-1 and apoF particles was approximately 1:2, corresponding roughly to the applied sample concentrations of 6 mg/mL and 10 mg/mL (**Fig. 6f**) and indicating a rather homogenous mixing of the two sample solutions. The cryo-EM structure of apoF in this region resolved to 1.87 Å (EMD-55025), showing that the on-grid mixing still does allow to reach semi-atomic resolution. The **Suppl. Video 2** demonstrates the sequential writing of Nrs-1 and apoF onto the same cryo-EM grid.

**Fig. 6.**
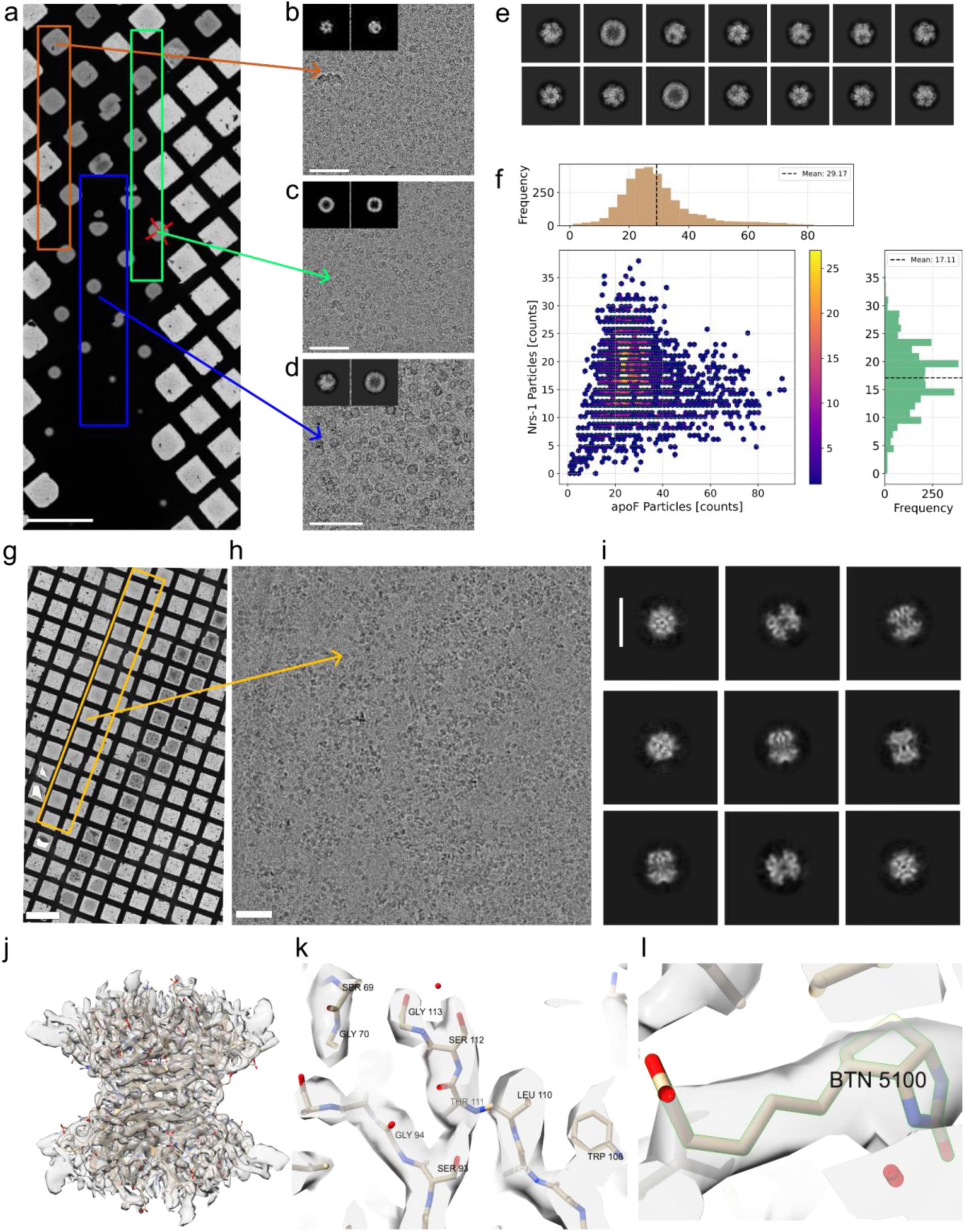
Writing and mixing of two samples on a cryoWriter and streptavidin-desthiobiotin interaction on a grid. **a** An overview of the vitrified grid showing the two different samples mixed within the grid (scale bar = 200 μm). Brown box: NrS-1 was applied, shown in **b** (scale bar = 80 nm). Green box: apoF protein was applied, shown in **c** (scale bar = 80 nm). Blue box: Mixing area, shown in **d** (scale bar = 60 nm). **e** 2D class averages from images acquired in the mixing region, show particles representing a population of Nrs-1 protein and apoF. **f** Statistical analysis of the NrS-1 and apoF particle distribution of the total 189,685 particles selected is shown in Fig. 5e. **g** Low magnification electron micrograph of streptavidin-desthiobiotin line writing shown in orange colored region (scale bar = 200 μm). **h** Electron micrograph of desthiobiotin-bound streptavidin molecules (scale bar = 100 nm). **i** 2D class averages (scale bar = 8 nm). **j** 3D reconstruction of desthiobiotin-bound streptavidin. **k** Representative densities of secondary structures, β-sheet. **l** Desthiobiotin density in the streptavidin binding pocket

### On-grid mixing: Streptavidin-desthiobiotin interaction

The cryowriter opens up the possibility of performing on grid biochemistry by mixing of protein and ligand or substrate directly on the grid and visualization of the binding results by cryo-EM. This opens up the possibility of trapping substrate intermediate states upon protein interaction for example in catalytic reactions. Current methods used to obtain time-resolved results mostly allow for an observation of a reaction time interval of 10-50 ms. Most of these methods, such as microfluidic mixing or electrospraying, rely on some form of sample mixing shortly before application to the grid. Even longer reaction times, in the order of seconds, can for example be achieved by manual mixing of the components before application to the grid ^27^. With the cryoWriter mixing in the time scale of the spot-to-plunge time, which is ∼200ms, is possible, closing the gap in available mixing times currently present. In addition to this, our approach allows for on-grid mixing of the samples, possibly enabling observation of multiple time points, or reactant gradients on the same grid.

To investigate these on-grid protein–ligand interactions, we performed two-line writing as described above, using streptavidin and desthiobiotin as model systems. Desthiobiotin was first deposited onto a cryo-EM grid at a concentration of 3 mM, using a 5.5 nL sample volume and a writing speed of 5 mm/s. A second line, spaced 300 µm from the first, was written using a 4.5 nL volume of streptavidin at a concentration of 3 mg/mL, also at a speed of 5 mm/s (**Fig. 6g**). The grid was plunge-frozen 200 ms after deposition. The **Suppl. Video 3** demonstrates the two-sample writing of biotin and streptavidin. We recorded 2,209 images in the zone between the two lines (**Fig. 6h,i**). From these 190,686 particles were initially extracted. After processing 47,564 particles contributed to the final reconstruction yielding a resolution of 2.97 Å (**Suppl. Table 4**). In the final map (**Fig. 6j**) secondary structures are clearly visible (**Fig. 6k**). The quality of this map allowed us to identify the desthiobiotin bound to streptavidin (**Fig. 6i**).Preparation of cryo-EM grids for streptavidin alone, however, resulted in particles showing a strong orientation bias when preparing grids with both, the cryoWriter as well as the TFS Vitrobot Mark IV (**Suppl. Fig. 7**).

## Discussion

Our results demonstrate the cryoWriter as a versatile grid preparation robot that can prepare high-quality cryo-EM grids from a few nanoliters of sample solution. In addition, the cryoWriter combines the grid preparation with an inverted light microscope for sample observations prior to freezing, a cooled sample storage box, a glow-discharger, and a versatile grid handling system that allows to prepare grids at dew point temperature for sample application, plunge-freeze the grid within 200 msec, and store the grids under liquid nitrogen.

We present the application of the cryoWriter to routinely freeze cryo-EM grids with various samples, including helical (TMV), single particle (apoF) and membrane protein particles in detergent (TRPM4). Grids can be plunged immediately after sample writing, or the plunging can be delayed for a few seconds prior to plunging, as also applied with other grid preparation robots ^6^. In our hands and with the samples tested, a delay time of around 3 seconds was optimal, with the cryoWriter operated at 70% relative humidity and the grid, launchpad, capillary and gripper tweezer all cooled to dew-point temperature.

Grids could be prepared from a few nanoliters of sample per grid, with samples at concentrations around 5 to 10 mg/mL. Variation in the spot-to-plunge time and multiple writing allowed to prepare grids with suitable particle density without the need of highly concentrated samples. Importantly, the microfluidic capillary of the cryoWriter was able to pick up sample volumes as small as 20 nanoliters and prepare a grid from that by applying 5 nanoliters onto the grid in a spiral or line writing pattern. Since handling of sample volumes of 20 nanoliters in Eppendorff tubes is not practical, a minimal sample volume of 3 microliters is usually required in order to prevent immediate drying of the sample when the Eppendroff cup is exposed to air during handling. After freezing several grids, each consuming only a few nL, the remaining volume from the 3µL sample could then still be used for other purposes. With these minimal volume requirements, the cryoWriter occupies a unique niche among the different cryo-EM grid preparation robots.

For the small protein streptavidin, preferred particle orientation on grids remained challenging. In such cases, usage of grids covered by a pristine layer of graphene that is nevertheless hydrophilic due to aromatic functionalization ^28^, or other treatments ^29,30^ could be helpful in reducing the preferred orientation issue, as these attract protein particles to the graphene water interface (GWI) and reduce the adsorption of particles at the air-water interface. These surface treatments were not tested here. However, we were able to utilize the cryoWriter to prepare grids from the NrS-1 sample, which was found to be much less affected by preferred orientation than when prepared with the other freezing methods tested. We processed the NrS-1 data for 3D reconstruction without applying any symmetry (C1 symmetry), in order to probe for non-symmetric particle deformations from air-water interface interactions. These were not observed.

Alternatively, processing the 6-fold symmetric particle dataset with C6 symmetry makes the particle distribution appear more uniform. While this can improve resolution for symmetric particles, as shown in **Suppl. Fig. 5**, it can also mask true orientation bias present in the dataset. At least for NrS-1, grid preparation with the cryoWriter was found to be sufficiently different from all other tested methods to result in a more random particle orientation, giving an additional option for this challenging sample.

Finally, we present an application of the cryoWriter, where two different samples were written onto the same grid. The sample particles diffused into each other, so that in the zone between the two written lines, a mixture of particles was imaged. This could also be used to perform on-grid incubation experiments, whereby the time between grid writing and plunging can be varied between 200 msec (the time required for grid plunging with the gripper) and a few seconds (not longer to prevent denaturation of the samples due to excessive draining or drying).

In conclusion, the cryoWriter is a versatile grid preparation robot that allows to prepare high-quality cryo-EM grids in a reproducible and consistent manner from nanoliter amounts of sample solution. The included light microscope, and the possibility to apply the sample or several samples to the grid in custom-programmable manners, such as double-writing the same sample twice, or writing two different samples near each other onto the grid, invite for a wide area of possible future applications of this instrument.

## Methods

### Protein purification

The tobacco mosaic virus (TMV) sample was a kind gift from Daniel Mann and Carsten Sachse (Ernst Ruska-Center for Microscopy and Spectroscopy with Electrons, ER-C, Wilhelm-Johnen-Straße, D-52428 Jülich). TMV was used at a concentration of 20 mg/mL for high-resolution cryo-EM imaging.

Horse spleen apoferritin was purchased from Sigma-Aldrich Chemie GmbH (product number A3660), dissolved in a buffer containing 25 mM HEPES-NaOH (pH 7.5), 150 mM NaCl and further purified by size exclusion chromatography on a pre-equilibrated Superose 6 gel filtration column. The peak fractions were concentrated on 100 K Amicon Ultra-15 concentrators to 10 mg/mL for single-particle cryo-EM.

The full-length *H. sapiens* TRPM4 was expressed and purified from HEK293F cells grown in suspension. For expression, HEK293F cells were transfected with 1 mg of a plasmid containing the TRPM4 gene per liter of cells using PEI (polyethylenimine). The cultures were grown at 37 °C and 5% CO_2_ for 48 h. The cultures were harvested by centrifugation at 3000 x g for 30 min at 4 °C and washed in 1x PBS, followed by another round of centrifugation. The pellets were carefully resuspended in lysis buffer containing 25 mM HEPES-NaOH (pH 7.5), 200 mM NaCl and supplemented with cOmplete™ EDTA-free Protease Inhibitor Cocktail (Roche). 4 tablets of protease inhibitor cocktail were added per 100 ml of buffer. Following resuspension, the cells were lysed by a single pass through a microfluidizer (Microfluidics™) at 15,000 psi and the membrane fraction was harvested by centrifugation using an Optima XPN-100 ultracentrifuge (Beckman Coulter) with the Ti45 rotor and spun at 70,560 × g for 30 min at 4 °C. The resulting pellets were stored at −80 °C. For detergent solubilization, 12 g of pellet was resuspended in 30 ml of solubilization buffer containing 25 mM HEPES-NaOH (pH 7.5), 200 mM NaCl, 1% Glyco-diosgenin (GDN, its critical micellar concentration is 0.002 to 0.003 %) (Avanti Pola lipids) supplemented with 2 tablets of cOmplete™ EDTA-free Protease Inhibitor Cocktail (Roche). The resuspended pellet was homogenized manually in a 40 ml Kimble glass homogenizer (Sigma). The tube containing the mixture was then placed in a bottle with a stir bar and left to incubate at 4 °C for 2 h. The homogenate was clarified by centrifugation for 30 min at 70,560 × *g* at 4 °C in an Optima XPN Ultracentrifuge (Beckman Coulter) using a Ti-45 rotor. The supernatant, which contains soluble FLAG-tagged HsTRPM4 mixed with 1 ml of Anti-FLAG**®** M2 affinity gel (Millipore, Billerica, MA) pre-equilibrated with wash buffer containing 25 mM HEPES-NaOH (pH 7.5), 200 mM NaCl and 0.02% GDN. The beads were washed with 100 mL of wash buffer containing 25 mM HEPES-NaOH (pH 7.5), 200 mM NaCl, and 0.02% GDN and eluted with 4 mL elution buffer containing 25 mM HEPES-NaOH (pH 7.5), 200 mM NaCl, 120 µg/ml of 3xFLAG peptide and 0.02% GDN. The purified protein was run on a Superose 6 gel filtration column pre-equilibrated with wash buffer and the peak fraction was concentrated on 100 K Amicon Ultra-15 concentrators (Millipore, Billerica, MA) to an absorbance at 280 nm of 6.0 to prepare cryo-EM grids.

Full-length bacteriophage NrS-1 DNA polymerase with codon optimized for *E. coli* expression was synthesized with a N terminal FLAG tag and cloned into pET28a vector (GenScript Biotech). NrS-1 DNA polymerase was expressed and purified from *E. coli* BL21(DE3). For expression of NrS-1 DNA polymerase chemically competent *E. coli* BL21(DE3) was transformed with the pET28a plasmid containing the kanamycin resistance gene for NrS-1 expression and grown overnight at 37 °C on Luria broth (LB) agar plates containing both selection antibiotics (50 μg ml^−1^ of kanamycin). The colonies obtained were streaked from the plate and transferred to 50 ml of 2xYT medium containing kanamycin and grown overnight at 37 °C with shaking. Then, 40 ml of the overnight culture was used to inoculate 4 l of 2xYT medium containing the selection antibiotics. The cultures were grown at 37 °C with shaking at 190 r.p.m. until they reached an absorbance at 600 nm of 0.5–0.7, then incubated on ice for 1 h, before inducing protein expression with 0.5 mM IPTG for 18 h at 20 °C. The overnight cultures were harvested by centrifugation at 3,000*g* for 30 min at 4 °C. The resulting supernatant was discarded and the pellet was resuspended in 100 ml of cold lysis buffer (25 mM HEPES-NaOH (pH 7.5), 500 mM NaCl, 10% glycerol, and 1 mM 2-mercaptoethanol) supplemented with two tablets of complete EDTA-free Protease Inhibitor Cocktail (Roche) and 5,000 units of Turbonuclease (Jena Bioscience) before lysis through a single pass in a Microfluidizer (Microfluidics™) at 15,000 psi. The lysate was clarified by centrifugation for 30 min at 70,560*g* and 4 °C in an Optima XPN Ultracentrifuge (Beckman Coulter) using a Ti-45 rotor. The supernatant contained soluble FLAG-tagged NrS-1, therefore purification was performed using anti-FLAG M2 affinity gel (Millipore): the beads were washed with 100 ml of wash buffer (25 mM Hepes-NaOH, pH 7.5, 150 mM NaCl, 10% glycerol and 1 mM 2-mercaptoethanol) and eluted with 4 ml of elution buffer (25 mM Hepes-NaOH, pH 7.5, 150 mM NaCl, 10% glycerol, 120 µg ml^−1^ of 3× FLAG peptide and 1 mM 2-mercaptoethanol), followed by buffer exchange and concentration on a 100K Amicon Ultra-15 concentrators (Millipore) and further purified by gel filtration chromatography on a 10/300 GL Superose 6 gel filtration column (Cytiva Life Sciences) in gel filtration buffer (25 mM Hepes-NaOH, pH7.5, 150 mM NaCl and 1 mM dithiothreitol (DTT)). Peak fractions (as determined by the chromatograms with ultraviolet light of 280 nm) generated from the Unicorn software (v.7.1) containing purified NrS-1 (as determined by sodium dodecylsulfate (SDS)–polyacrylamide gel electrophoresis (PAGE) analysis), were pooled and concentrated to a final concentration of 6 mg/ml to prepare cryo-EM grids.

Streptavidin was purchased from BioConcept AG (product number N7021S) at a stock concentration of 1 mg/mL. For grid preparation, the protein was concentrated to a final concentration of 3 mg/mL in buffer containing 25 mM HEPES-NaOH (pH 7.5) and 75 mM NaCl. d-Desthiobiotin (Sigma) was resuspended in the same buffer to a concentration of 3mM to be used for cryo-EM experiments.

### Sample preparation

Quantifoil R1.2/1.3 Cu 300 grids (Quantifoil Micro Tools, GmbH, Germany) were used as sample carriers for TMV, apoferritin, and TRPM4 samples. These are copper grids that are covered with a carbon film that has holes of 1.2 micrometer diameter, separated from each other by a surrounding carbon film of 1.3 µm spacing. For Streptavidin-desthiobiotin, Quantifoil^®^ Active grids composed of 300-mesh copper with nanowires, and holey carbon support films featuring a round-hole geometry (1.2 µm diameter, 0.8 µm edge-to-edge spacing) were used. The grids were rendered hydrophilic by glow-discharging for 60 seconds in low-pressure air, using the on-board glow discharge system in the cryoWriter for apoferritin grid preparation. Alternatively, grids were glow-discharged using a PELCO easiGlow™ system at 15 mA for 60 seconds. Samples were vitrified within 200 ms after sample writing or with more delay, by plunging into liquid ethane, cooled to -180°C with the help of liquid nitrogen and the cryoWriter’s thermostat, before automatically transferring the grids to the cryo-EM grid storage box under liquid nitrogen.

### Single-Particle Cryo-EM Data Collection

Cryo-EM images were recorded with a Thermo Scientific 300 kV Titan Krios G4 equipped with a Falcon 4i detector (TFS). Data were acquired using the automated data collection software EPU (TFS). For TMV, data were collected at a magnification of 270kx and a pixel size of 0.46 Å at the specimen level, 80 e^−^/Å^2^ total electron dose, and a defocus between −0.35 and −0.85 μm, recording dose-fractionated data as electron event recordings (EER). For apoferritin, data were collected at a magnification of 120kx and a pixel size of 0.66 Å, 60 e^−^/Å^2^ total electron dose, and defocus values between −0.5 and −2.0 μm. For TRPM4, data were collected at a magnification of 165kx, a pixel size of 0.73 Å, 50 e^−^/Å^2^ total electron dose, and defocus values between −0.35 and −2.0 μm. For streptavidin, data were collected at a magnification of 165kx, a pixel size of 0.732 Å, a total electron dose of 60 e^−^/Å^2^, and defocus values between −1 and −2 μm. All image data were stored as electron event representation (EER) (**Suppl. Table 3**).

### Image processing

Cryo-EM raw EER files were imported and processed in CryoSPARC ^31^. Motion correction was performed using Patch Motion Correction, followed by Patch CTF Estimation. After CTF correction, only micrographs with an estimated resolution better than 6 Å were selected for further processing. For TMV, a total of 5,824 movies were collected. Following CTF estimation, the best 3,732 micrographs were retained for further analysis. Filament tracing was performed in CryoSPARC, and 484,223 helical segments were picked and extracted with a box size of 512 pixels. Three rounds of 2D classification were conducted to select high-quality class averages. Helical symmetry was subsequently applied for 3D reconstruction. For apoferritin, 6,121 high-quality micrographs were used for processing. The Blob Picker tool was employed for initial particle picking and template generation, followed by a second round of template-based picking. A total of 631,925 particles were extracted using a 448-pixel box size. The dataset underwent three rounds of 2D classification for quality screening, resulting in 489,099 particles being retained for 3D reconstruction with O symmetry imposed. For TRPM4, 3,535 micrographs were collected, of which 3,458 passed CTF-based quality screening. A total of 695,101 particles were extracted with a box size of 512 pixels and Fourier-cropped to 440 pixels. Following three rounds of 2D classification and removal of low-quality particles, 69,878 particles were selected for 3D reconstruction with C4 symmetry applied. For the streptavidin–desthiobiotin complex, a total of 2,209 movies were collected, of which 1,934 were retained following CTF-based quality screening. An initial set of 1,018,875 particles was extracted with a box size of 128 pixels and Fourier-cropped to 64 pixels. After three rounds of 2D classification and removal of low-quality particles, the remaining particles were re-extracted at 448 pixels and Fourier-cropped to 256 pixels. A final subset of 47,564 high-quality particles was selected for 3D reconstruction, with D2 symmetry imposed. Single-particle data collection and map statistics are summarized in **Suppl. Table 3**.

### Model refinement

The atomic models for TMV (6RLP ^32^), apoferritin (6PXM ^33^), TRPM4 (8RCR ^15^), and streptavidin (6J6J ^34^) were manually superimposed onto the sharpened cryo-EM maps in ChimeraX ^35^. The models were iteratively refined, including all-atom and chain refinement in Coot ^36^, followed by real-space refinement using Phenix ^37^, and final optimization to ensure the best fit to the maps. The refined models were then used to generate graphical figures in ChimeraX.

## Data availability

Raw images are available at EMPIAR under accession numbers XXXX.

Reconstructed 3D volumes are available at the EMDB under accession numbers EMD-54957, EMD-54970, EMD-55006, EMD-54984, EMD-55000, EMD-55025, and EMD-55027

## Acknowledgments

We thank Prof. Andreas Engel and his colleagues at cryoWrite AG, Basel, Switzerland, for excellent support and fruitful discussions. Cryo-EM data were partially acquired at the Dubochet Center for Imaging (DCI) in Lausanne, Switzerland. We acknowledge E. Uchikawa, B. Beckert, S. Nazarov, and A. Myasnikov at DCI-Lausanne for their assistance.

This work was in part supported by the Swiss National Science Foundation (grant 200021_200628), and by the European Union (ERC 4D-BioSTEM, No 101118656). Views and opinions expressed are, however, those of the authors only and do not necessarily reflect those of the European Union or the European Research Council Executive Agency. Neither the European Union nor the granting authority can be held responsible for them.

## Author contributions

HS conceived and managed the project. KVC designed and conducted the experiments, KVC and MK analyzed the results. BE purified protein samples. IM and KVC collected the single-particle cryo-EM data. MDF performed an early calibration experiment. JR supported cryo-EM data collection. KVC performed the cryo-EM data processing and protein structure analysis. BE and IM assisted by reviewing the cryo-EM data processing. KVC and MK wrote the initial draft of the manuscript. KVC, MK, and HS revised and edited the manuscript, with contributions from all authors.

## Additional information

The Supplementary Information contains:

● Suppl. Fig. S1: Resolution analysis of cryo-EM reconstructions by Fourier Shell Correlation (FSC).
● Suppl. Fig. S2: High-resolution single particle reconstruction of apoF with line writing prepared using the cryoWriter.
● Suppl. Fig. S3: Atlas images of grids, prepared with different writing patterns.
● Suppl. Fig. S4: Ice thickness measurement of cryoWriter grid using cryo-ET from Position 2 to Position 6.
● Suppl. Fig. S5: Analysis of the NrS-1 particle orientation in cryo-EM grids.
● Suppl. Fig. S6: Scheme of writing two different samples onto the same cryo-EM grid.
● Suppl. Fig. S7: Overview and 2D classification of preferential orientation of streptavidin protein.
● Suppl Text 1: Calculation of the expected particle numbers in an image
● Suppl Text 2: Calculation of expected trace thickness.
● Suppl. Table 1: Particle density in cryo-EM grids prepared with different waiting times between writing and plunge-freezing.
● Suppl. Table 2: Electron micrographs showing particle distribution with varying concentrations.
● Suppl. Table 3: Statistical parameters of the determined protein structures from cryo-EM grids prepared with the CryoWriter.
● Suppl. Video 1: The cryoWriter in operation
● Suppl. Video 2: Spiral writing
● Suppl. Video 3: Demonstration of the writing of two separate samples onto the same grid.

## Supplementary Information

**Fig. S1.**
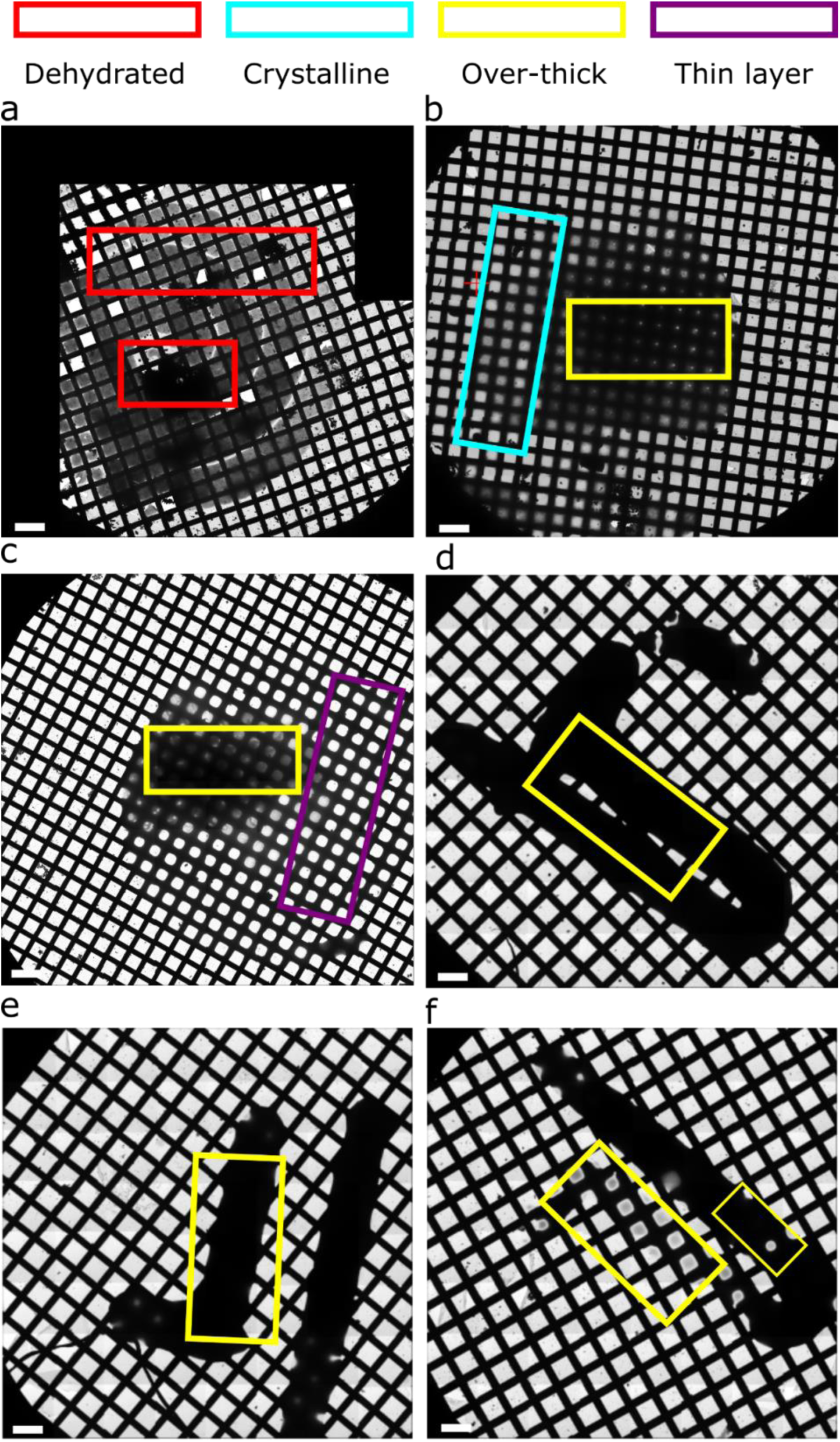
Atlas images of grids, prepared with different writing patterns. **a** Overview image of the spiral writing pattern of the dehydrated apoF sample caused by evaporation before freezing. **b** Atlas image of overly-thick ice at the center and crystalline ice. **c** Combination of both, overly-thick and thin ice. **d-e** Line writing of the apoF sample leads to the formation of overly-thick ice (scale bar = 200 µm).

**Fig. S2.**
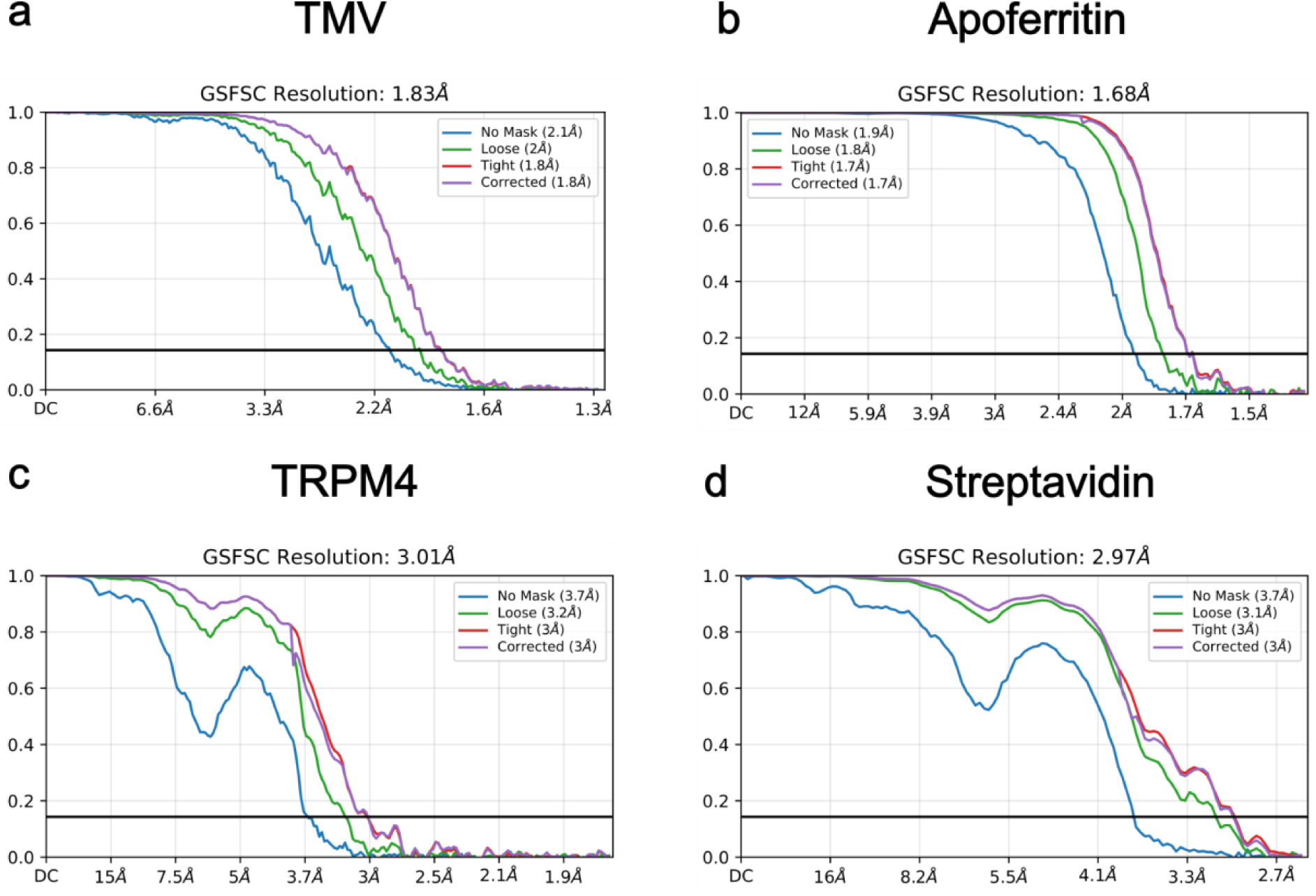
Resolution analysis of cryo-EM reconstructions by Fourier Shell Correlation (FSC). **a** TMV at 1.83 Å resolution. **b** apoF at 1.68 Å resolution. **c** TRPM4 at 3.01 Å resolution. **d** Streptavidin and desthiobiotin complex at 2.97 Å resolution.

**Fig. S3.**
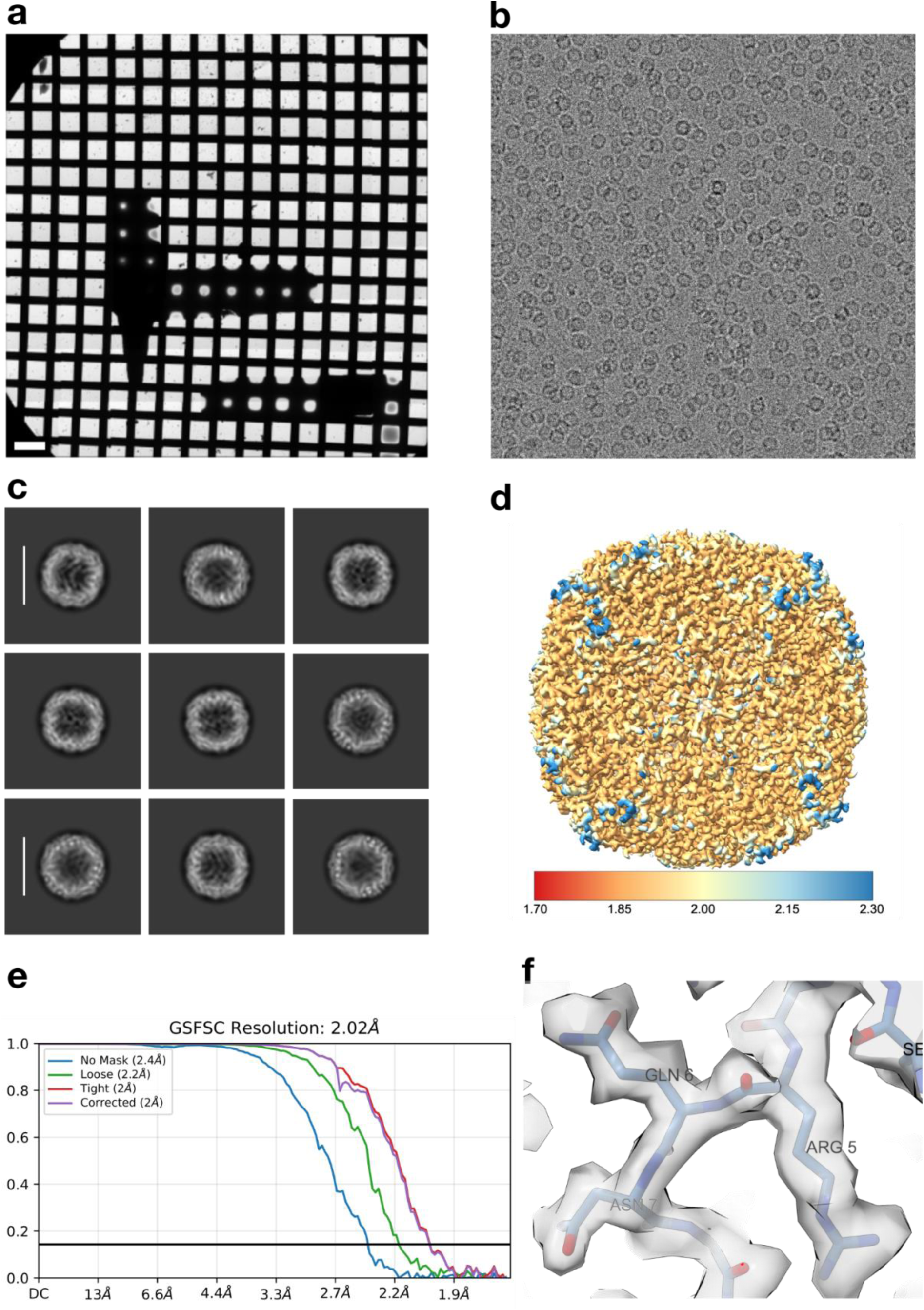
High-resolution single particle reconstruction of apoF with line writing prepared using the cryoWriter. **a** Overview of the vitrified grid (scale bar = 200 μm). **b** Representative cryo-EM micrograph of apoF (Scale bar = 50 nm). **c** 2D class average of apoF (scale bar = 11 nm). **d** Local resolution map colored according to the assigned scale (scale in Å). **e** FSC curve showing the resolution at 2.02 Å. **f** An exemplary image showing the cryo-EM densities of apoF fit well into the atomic model.

**Fig. S4.**
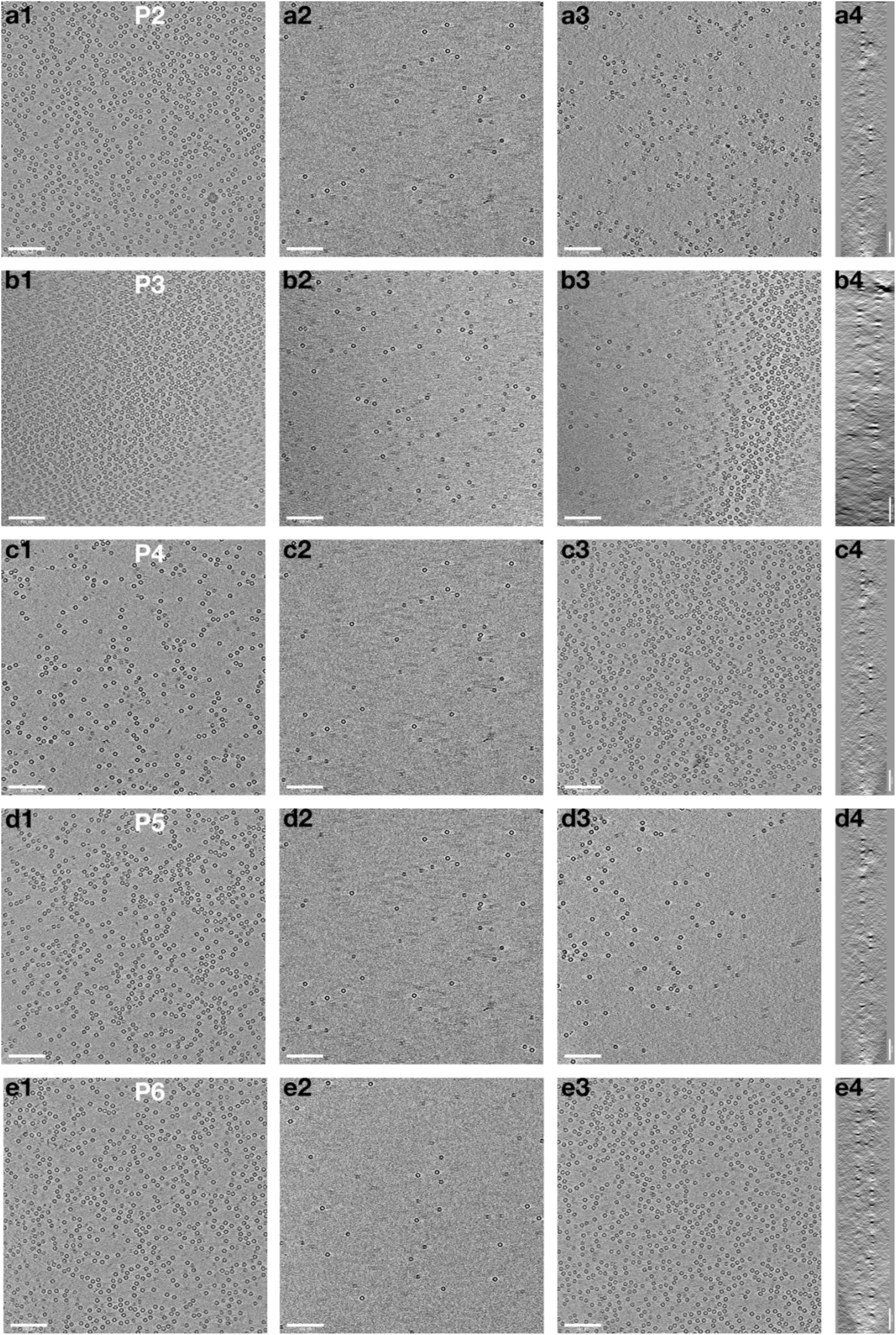
Measurements of ice thickness and particle distribution in cryoWriter grids. a-e. Panels from top to bottom are from positions 2 to 6 on the ApoF grid shown in Fig. 3a. Sub-panels 1 to 4 from left to right depict **1:** Tomographic reconstruction, showing the top layer of the 3D reconstruction. **2**: showing the central layer, and **3**: the bottom layer of the 3D reconstruction. **4**: The side-view of the 3D reconstruction allows to recognize the 3D distribution of particles to the two surface layers of the ice. (scale bars = 100 nm).

**Fig. S5.**
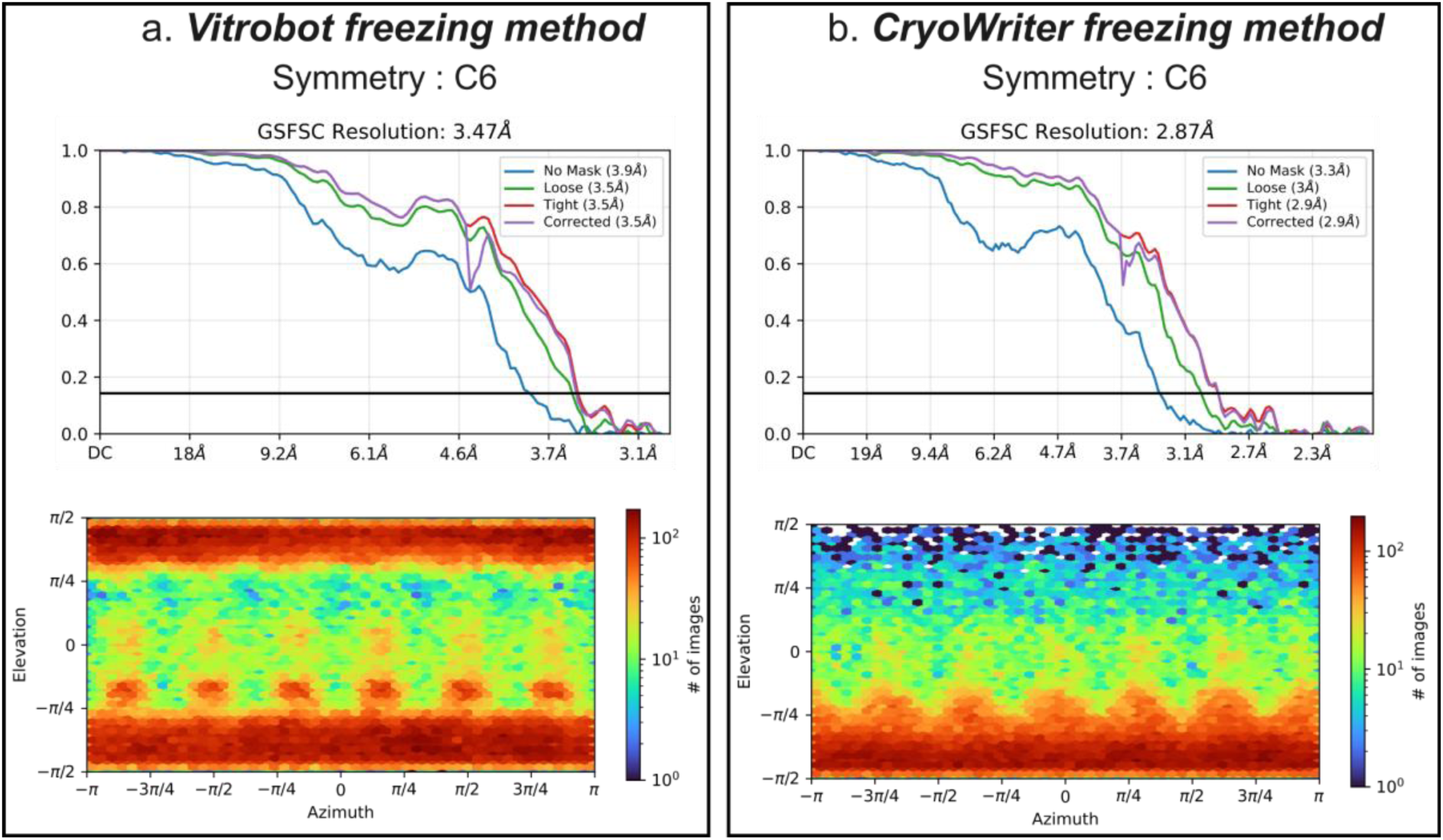
Analysis of the NrS-1 particle orientation in cryo-EM grids. **a** Vitrobot freezing method with C6 symmetry applied, corresponding FSC curve at 3.47 A resolution, and a particle viewing direction distribution for 133,779 particles. The resolution improved when compared to C1 symmetry. **b** CryoWriter freezing method when applied to C6 symmetry, corresponding FSC curve showing 2.87 Å resolution and particle viewing direction distribution with 96,862 particles. The resolution improved when compared to C1 symmetry.

**Fig. S6.**
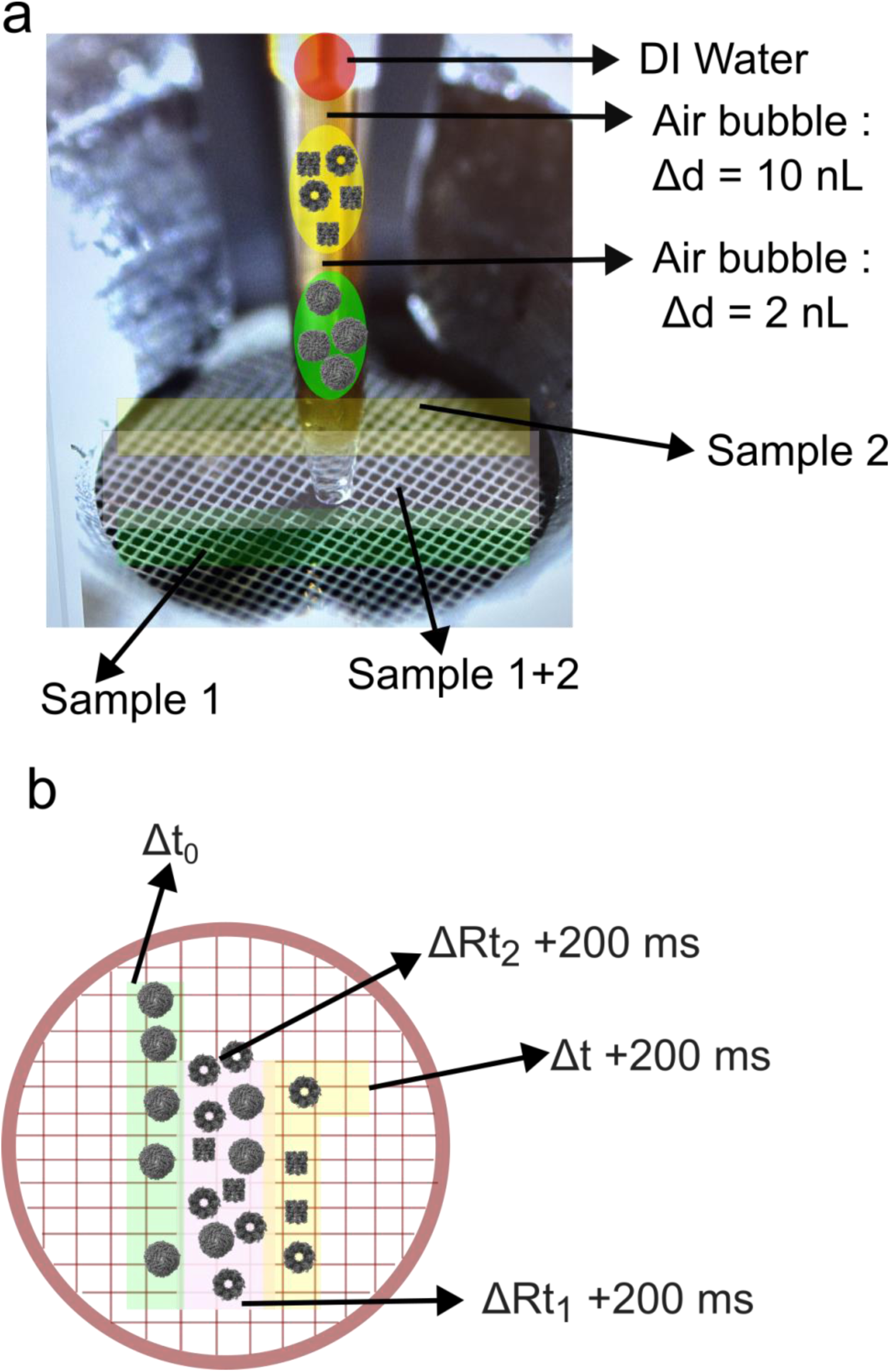
Scheme of writing two different samples onto the same cryo-EM grid. **a** Schematic diagram illustrating these two different sample writing strategies on the same grid. **b** Schematic diagram illustrating time resolution approximation by writing two different samples on a cryo-EM grid.

**Fig. S7.**
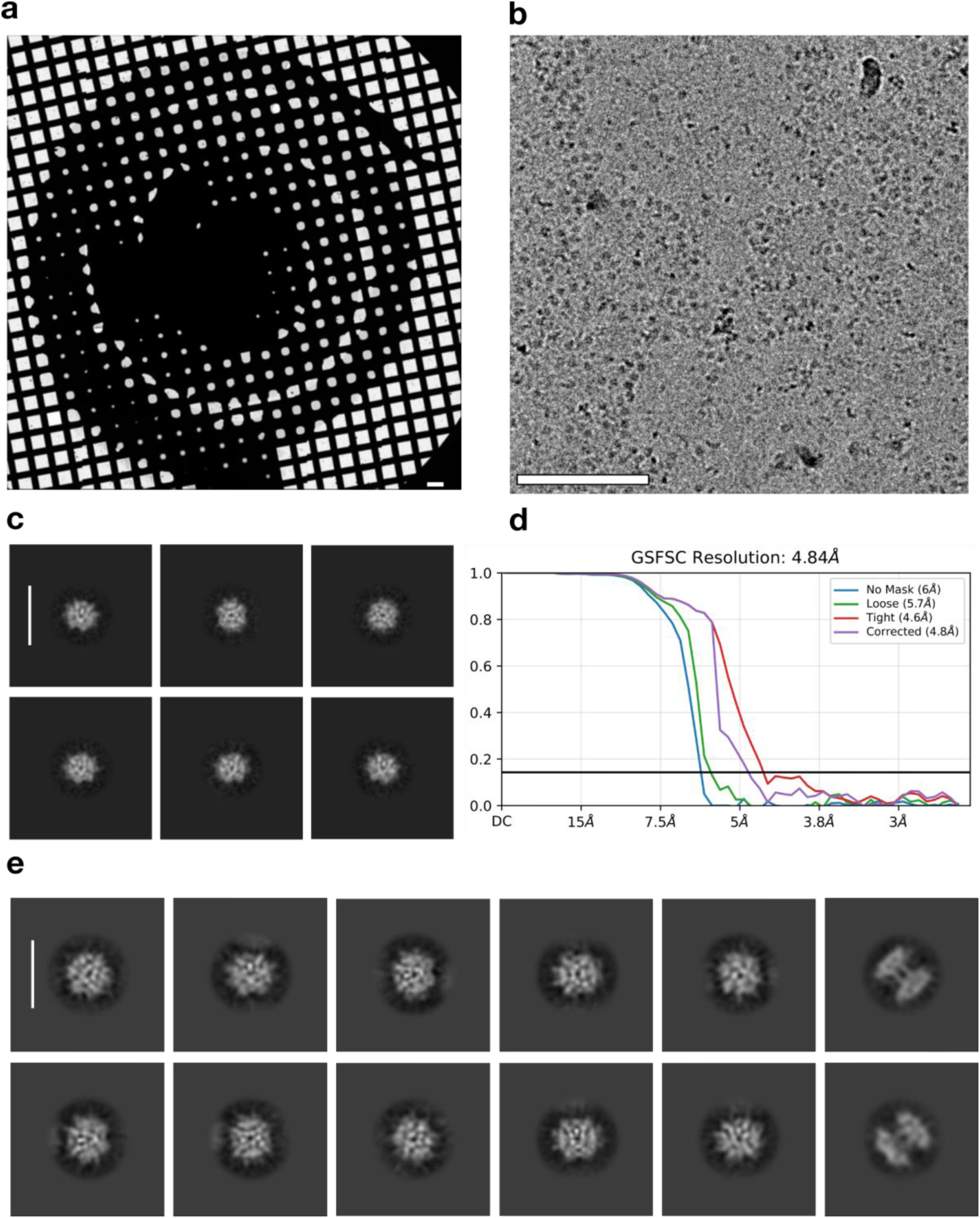
Overview and 2D classification of preferential orientation of streptavidin protein. **a** Overview of the vitrified grid of streptavidin (scale bar = 200 μm). **b** Representative cryo-EM micrograph of the streptavidin protein alone (Scale bar = 50 nm). **c** 2D class average of streptavidin (scale bar = 9 nm). **d** FSC curve shows the resolution is at 4.84 Å due to strong preferential orientation. **e** 2D class averages of streptavidin showing preferential orientation frozen with vitrobot (Scale bar = 7 nm).

### Suppl. Text 1: CryoWriter freezing workflow

1. Start by filling the cryopot with liquid nitrogen, followed by filling the ethane pot with liquification ethane, and maintaining the ethane temperature between 180 and 183 °C.
2. The eppendorf tube containing the sample is placed in a nano-incubator, with the temperature adjusted according to experimental requirements. In routine operation, the nano-incubator is maintained between 4 °C and 8 °C.
3. The relative humidity control is then activated, and dew point offset temperatures are set for the tweezer, capillary, and launchpad. These values are user-defined; typical operating conditions are 60-70% relative humidity and a dew point temperature offset of +2 °C.
4. Grids are subsequently loaded into the grid loading station. Using the gripper/tweezer, each grid is automatically transferred to the glow discharge station to render the surface hydrophilic. Glow discharge parameters, such as duration and discharge current, are user-specified. Alternatively, grids may be pre-glow-discharged outside the cryoWriter system before loading.
5. Following glow discharge, the tweezer automatically transfers the grid to the launchpad, which is maintained at the dew point temperature, typically 16-18 °C.
6. The capillary is then positioned in the nano-incubator to aspirate the desired volume of sample, with both volume and aspiration speed determined by the user. The capillary is subsequently moved back to the launchpad, where the grid is positioned for sample deposition.
7. Sample application parameters, including writing pattern (e.g., spiral or linear), writing speed, deposition rate, and spacing between writing are user-controlled.
8. Finally, the grid is plunge-frozen into liquid ethane, after which it is automatically transferred into a storage puck.

### Suppl. Text 2: Calculation of the expected particle numbers in an image

#### Calculation of protein particles per field of view in an electron microscope

Apoferritin is a spherical particle with 24-fold symmetry. The particles have a molecular weight 𝑤 of 483 kDa or 483,000 g/mol, and we assume a mass concentration 𝑐 of 10 mg/mL or 10 g/L. The mass concentration can be expressed as particle concentration with:

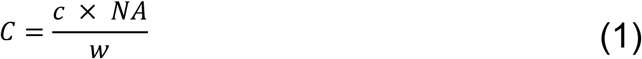

with 𝑁𝐴 = 6*․*022 × 10^23^ particles/mol being Avogadro’s number. The apoferritin sample therefore was available at a concentration of

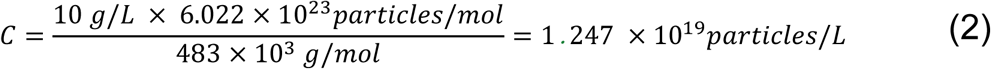

or 1 *pL* = 12.47 × 10^6^ *particles*

Let us assume an ice thickness of 100 nm. The sample volume within a square micrometer is then 0.1 µm^3^. Using the fact that 1 µm^3^ = 1 × 10^-3^ pL, we note that in a square micrometer we should encounter a sample volume of 1 × 10^-4^ pL. That volume in the above example would contain 1247 particles, or result in a particle density of 1247 particles per square micrometer.

### Suppl. Text 3: Calculation of expected trace thickness

#### Variable definition

𝜔 = angular velocity [rad/s]

𝜃 = azimuth angle of of the spiral [rad]

𝑡 = time [s]

r = radius of the spiral [m]

𝑎_𝑐_ = acceleration of the capillary during writing [m/s^2^]

𝑣 = speed of the capillary on the sample [m/s]

𝑄 = flow rate of the sample [L/s]

𝑉 = dispensed sample amount [L] = 𝐴 × ℎ

𝐴 = surface area covered by sample [m^2^]

ℎ = initial height of the written sample trace on the grid [m]

𝑑 = initial diameter of the written sample trace on the grid [m]

#### Capillary writing at constant angular velocity and constant dispensing rate

When writing the sample onto the grid in a spiral pattern at a constant angular velocity (𝜔 = 𝑐𝑜𝑛𝑠𝑡.), leading to a constantly increasing azimuth angle (𝜃(𝑡) = 𝜔 × 𝑡, with 𝑡 being time), and using a slowly increasing radius (𝑟(𝑡)), a curved path is formed during which the capillary is moving at increasing speed when writing from the center to the outside of the spiral:

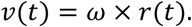

The capillary is exposed to an acceleration 𝑎_𝑐_(𝑡) as:

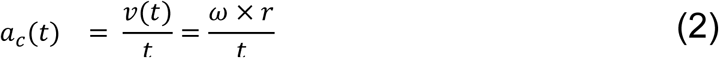

Assuming a constant width 𝑑 of the written trace on the grid, the covered area on the grid per time is:

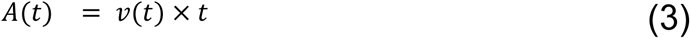

If the sample is dispensed at a constant rate (𝑄), e.g., 𝑄 = 1 𝑛𝐿/𝑠, then the amount (𝑉) of dispensed sample after a certain time is 𝑉(𝑡) = 𝑄 × 𝑡. The height before any evaporation of the written trace is ℎ(𝑡) = 𝛥𝑉(𝑡)/𝛥𝐴(𝑡).

The acceleration of the writing capillary 𝑎_𝑐_ then leads to a decrease of the initial trace height ℎ as:

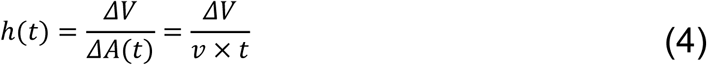

This shows that writing at constant angular velocity and constant dispensing rate leads to the generation of a spiral pattern that starts in the center with slower movements and finishes towards the outside with faster capillary movements, resulting in a decreasing sample thickness towards the outer areas of the spiral. This effect can be partly compensated with controlled drying, which would affect the first-written center of the spiral more than the outer areas of the spiral.

**Suppl. Table 1:**
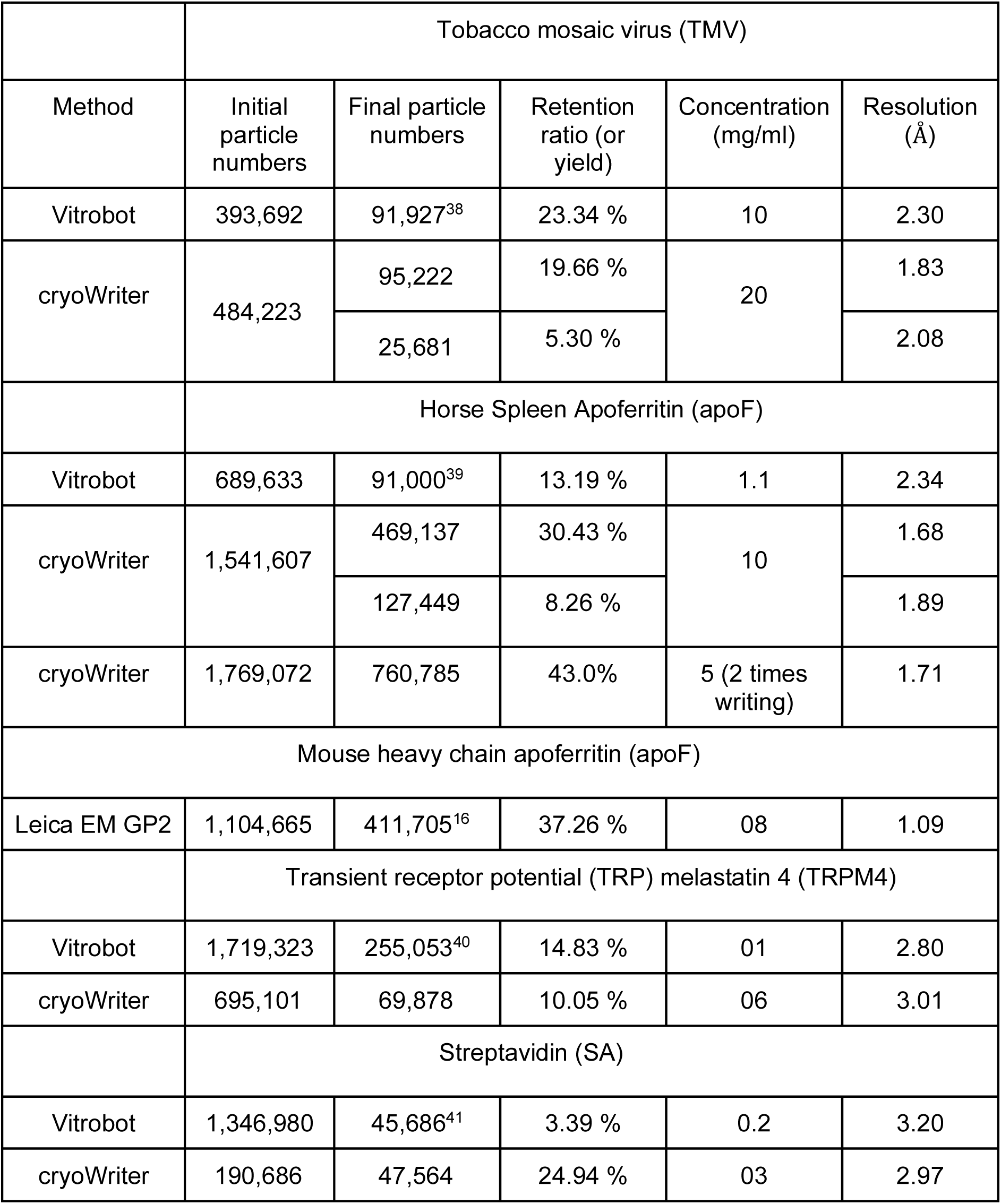
Quantitative comparison of particle yield for final 3D reconstruction and concentration between cryoWriter and conventional plunge freezing.

**Suppl. Table 2:** Particle density in cryo-EM grids prepared with different waiting times between writing and plunge-freezing. ApoF protein sample at a concentration of 5 mg/mL was written twice onto the same cryo-EM grid, followed by different waiting times before plunge-freezing. The images were recorded at strong defocus (i.e., high image contrast) on thin ice layers, so that all particles were strongly visible.

| Sample | Apoferritin |  |  |  |
| --- | --- | --- | --- | --- |
| Molecular weight | 483 kDa |  |  |  |
| Concentration [mg/mL] | 5 |  |  |  |
| Writing type | Spiral |  |  |  |
| Number of writings | 2 |  |  |  |
| Spot-to-plunge time [s] | 2 | 3 | 4 | 5 |
| Micrographs | 2 | 9 | 8 | 4 |
| Pixel size [Å/px] | 0.83 |  |  |  |
| Expected number of particles per $\mu\text{m}^2$ when assuming deposition of a 100nm layer of sample | 1247<br>(2 x 623 due to double-writing) | | | |
| Observed number of particles per $\mu\text{m}^2$ | 2,674 | 3815 | 5,537 | 3,071 |

**Suppl. Table 3:** Electron micrographs showing particle distribution with varying concentrations. The images were recorded at lower defocus (i.e., optimized for high resolution data collection, while providing lower contrast) on slightly thicker ice layers and in the vicinity of the carbon film. Only isolated particles suitable for high-resolution structural analysis were picked. This resulted in a lower particle count than in Suppl. Table 1.

| Sample | Apoferritin |  |  |  |
| --- | --- | --- | --- | --- |
| Molecular weight | 483 kDa |  |  |  |
| Concentration (mg/mL) | 2 | 5 | 5 | 10 |
| Writing type | Spiral | Spiral | Spiral | Spiral |
| Number of writings | 1 | 1 | 2 | 1 |
| Ice thickness assumption [nm] | 100 |  |  |  |
| Micrographs | 12 | 5 | 16,692 | 6,017 |
| Pixel size [Å/px] | 0.83 | 0.9 | 0.41 | 0.66 |
| Spot-to-plunge time [s] | 0.2 | 0.2 | 3 | 0.2 |
| Expected number of particles per $\mu\text{m}^2$ when assuming deposition of a 100nm layer of sample | 249 | 623 | 1,247 | 1,247 |
| Observed number of particles per $\mu\text{m}^2$ | 25 | 250 | 1631 | 1505 |
| Resolution | – | – | 1.71 | 1.68 |
| EMDB ID | – | – | EMD-55027 | EMD-54957 |

**Suppl. Table 4:** Statistical parameters of the determined protein structures from cryo-EM grids prepared with the cryoWriter.

| Sample | TMV | Apoferritin |  | TRPM4 | Strept-<br>avidin |
| --- | --- | --- | --- | --- | --- |
| Microscope | TFS Titan Krios G4, CFEG, Falcon 4i |  |  |  |  |
| Voltage [kV] | 300 |  |  |  |  |
| Grid type | Quantifoil R 1.2/1.3 on Cu 300 |  |  |  | Quantifoil<br>Active |
| Energy Filter (10 eV) | Selectris X | – | – | – | Selectris X |
| Sample Conc. [mg/mL] | 20 | 10 | 10 | 6 | 3 |
| Writing type | Spiral | Spiral | Line | Spiral | Line |
| Capillary diameter [μm] | 125 | 125 | 125 | 200 | 125 |
| Nominal Magnification | 270,000 | 120,000 | 96,000 | 165,000 | 165,000 |
| Electron exposure [e<sup>-</sup>/Å<sup>2</sup>] | 80 | 60 | 40 | 50 | 60 |
| Defocus [μm] | 0.35 to 0.85 | 0.5 to 2.0 | 0.5 to 2.0 | 0.35 to 0.85 | 1.0 to 2.0 |
| Pixel size [Å] | 0.46 | 0.66 | 0.83 | 0.732 | 0.732 |
| Micrographs | 5,824 | 6,121 | 4,923 | 3,535 | 2,209 |
| Initial particle numbers | 484,223 | 1,541,607 | 802,322 | 695,101 | 190,686 |
| Extraction Box Size | 512 | 448 | 320 | 440 | 256 |
| Particles per micrograph | 47 | 105 | 111 | 201 | 99 |
| Final particle numbers | 95,222 | 469,137 | 230,450 | 69,878 | 47,564 |
| Symmetry imposed | Helical | O | O | C4 | D2 |
| Guinier Plot B-factor [Å<sup>2</sup>] | 18.7 | 51.7 | 65.8 | 56.7 | 105 |
| FSC threshold | 0.143 |  |  |  |  |
| Map resolution [Å] | 1.83 | 1.68 | 2.02 | 3.01 | 2.97 |
| EMPIAR ID | TBD | TBD | TBD | TBD | TBD |
| EMDB ID | EMD-55006 | EMD-54957 | EMD-54970 | EMD-54984 | EMD-55000 |
| Fit PDB ID | 6RLP | 6PXM | 6PXM | 8RCR | 6J6J |

